# A prion accelerates proliferation at the expense of lifespan

**DOI:** 10.1101/2020.07.10.196584

**Authors:** David M. Garcia, Edgar A. Campbell, Christopher M. Jakobson, Mitsuhiro Tsuchiya, Acadia DiNardo, Matt Kaeberlein, Daniel F. Jarosz

## Abstract

Organisms often commit to one of two strategies: living fast and dying young or living slow and dying old. In fluctuating environments, however, switching between these two strategies could be advantageous. Lifespan is often inversely correlated with cell size and proliferation, which are both limited by protein synthesis. Here we report that a highly conserved RNA-modifying enzyme, the pseudouridine synthase Pus4/TruB, can act as a prion, endowing yeast with greater proliferation rates at the cost of a shortened lifespan. Cells harboring the prion can grow larger and exhibit altered protein synthesis. This epigenetic state, [*BIG*^+^] (<u>b</u>etter in <u>g</u>rowth), allows cells to heritably yet reversibly alter their translational program, leading to the differential expression of hundreds of proteins, including many that regulate proliferation and aging. Our data reveal a functional role for aggregation of RNA-modifying enzymes in driving heritable epigenetic states that transform cell growth and survival.

## Introduction

Cell size and proliferation are fundamental determinants of development, survival, and disease (Su and O’Farrell, 1998). Although these features can be independently controlled, coupling is common due to their dependence on the same biochemical building blocks (Su and O’Farrell, 1998; Turner et al., 2012). Growth and proliferation also impact lifespan across eukaryotes (Fontana et al., 2010; Kenyon, 2010; Pitt and Kaeberlein, 2015). The importance of tightly controlling each of these properties is underscored by the many mutations affecting growth and proliferation that are pathogenic, leading to cancer, developmental abnormalities, and myriad diseases of age (reviewed in (Hanahan and Weinberg, 2011; Saxton and Sabatini, 2017).

Organisms commonly alter the relationship between cell growth and proliferation during the course of development (Su and O’Farrell, 1998). In oocytes of *Drosophila melanogaster*, for example, a massive expansion in cytoplasmic volume without cellular division precedes a later step of numerous cellular divisions without cytoplasmic growth. The rates of cell growth and proliferation can be strongly coupled to nutrient sensing and to changes in metabolism (Efeyan et al., 2015; Turner et al., 2012). Thus, organisms can use genetically encoded signaling pathways to commit to different strategies depending on both their needs and their environment (Ivanov et al., 2015; Jung et al., 2018). Epigenetic tuning of cell growth and proliferation could in principle provide a stable yet reversible mechanism to alter the relationship between growth and proliferation according to differing needs in fluctuating environments. Histone modifications can enable such adaptation in a way that can be heritable over several mitotic divisions. However, apart from a few notable exceptions (Catania et al., 2020; Grewal and Klar, 1996; Nakayama et al., 2000), the majority of studied examples are erased during meiosis (Heard and Martienssen, 2014; Moazed, 2011).

Prions are a distinct class of epigenetic mechanism that can be faithfully transmitted through both mitotic and meiotic cell divisions (Brown and Lindquist, 2009; Cox, 1965; Garcia and Jarosz, 2014; Wickner, 1994). The unusual folding landscape of prion proteins, which allows the recruitment of proteins from the naïve to the prion fold, promotes a mode of inheritance that is both stable and reversible (Chakrabortee et al., 2016a; McKinley et al., 1983). For example, transient perturbations in molecular chaperone activity (Brown and Lindquist, 2009; Chernoff et al., 1995), specific environmental stressors (Garcia et al., 2016; Singh et al., 1979; Tuite et al., 1981), or regulated proteolysis (Ali et al., 2014; Kabani et al., 2014) can induce or eliminate prion states. It is now appreciated that this form of information transfer is far more common than previously realized (Chakrabortee et al., 2016b; Halfmann et al., 2012; Yuan and Hochschild, 2017).

Here we investigate the inheritance of a prion-based epigenetic state that alters yeast cell physiology, potentiating a tradeoff between proliferation and lifespan. We first describe the [*BIG*^+^] prion—driven by the conserved pseudouridine synthase Pus4 (known as TruB in mammals and bacteria)—which increases proliferation but shortens lifespan. We then quantitatively model the adaptive value of this *‘live fast, die young’* growth strategy in fluctuating environments. [*BIG*^+^] cells are larger and exhibit increased protein synthesis, as well as increased pseudouridylation activity. Finally, we find evidence for analogous epigenetic control of cell size in wild yeast populations. The epigenetic inheritance of an altered form of an RNA-modifying enzyme over long biological timescales, as occurs in [*BIG*^+^], thus provides a mechanism through which short-lived epigenetic modifications of nucleic acid can be perpetuated across generations.

## RESULTS

### Cells bearing a prion-like epigenetic element live fast and die young

We recently discovered more than 40 protein-based epigenetic elements in *Saccharomyces cerevisiae* that are both heritable and reversible upon transient perturbation of protein chaperone function (Chakrabortee et al., 2016a). Many of the proteins that underlie this behavior have the potential to regulate growth. One of these epigenetic states was induced by transient overexpression of the highly conserved pseudouridine synthase *PUS4*/TRUB, which catalyzes formation of a ubiquitous pseudouridine on U55 in tRNAs in bacteria, yeast and humans (Becker et al., 1997; Gutgsell et al., 2000; Zucchini et al., 2003). Mutation of U55 leads to large fitness deficits, second only to mutations in the tRNA anticodon loop, highlighting the functional importance of this nucleotide (Li et al., 2016). Mutation of U55 is also linked to deafness and diabetes in humans (Wang et al., 2016). We originally discovered this epigenetic state because transient overexpression of *PUS4* led to an enduring and heritable growth improvement in medium containing zinc sulfate (Chakrabortee et al., 2016a). Upon closer examination, we noticed that cells harboring the Pus4-induced element also achieved up to a ∼60% faster maximal proliferation rate than naïve cells in standard rich medium (YPD; **Fig. 1A**, p=0.0095, unpaired t-test). We therefore initially named this mitotically heritable element “Big^+”^, for <u>b</u>etter in <u>g</u>rowth, on the basis of its phenotype.

**Figure 1.**
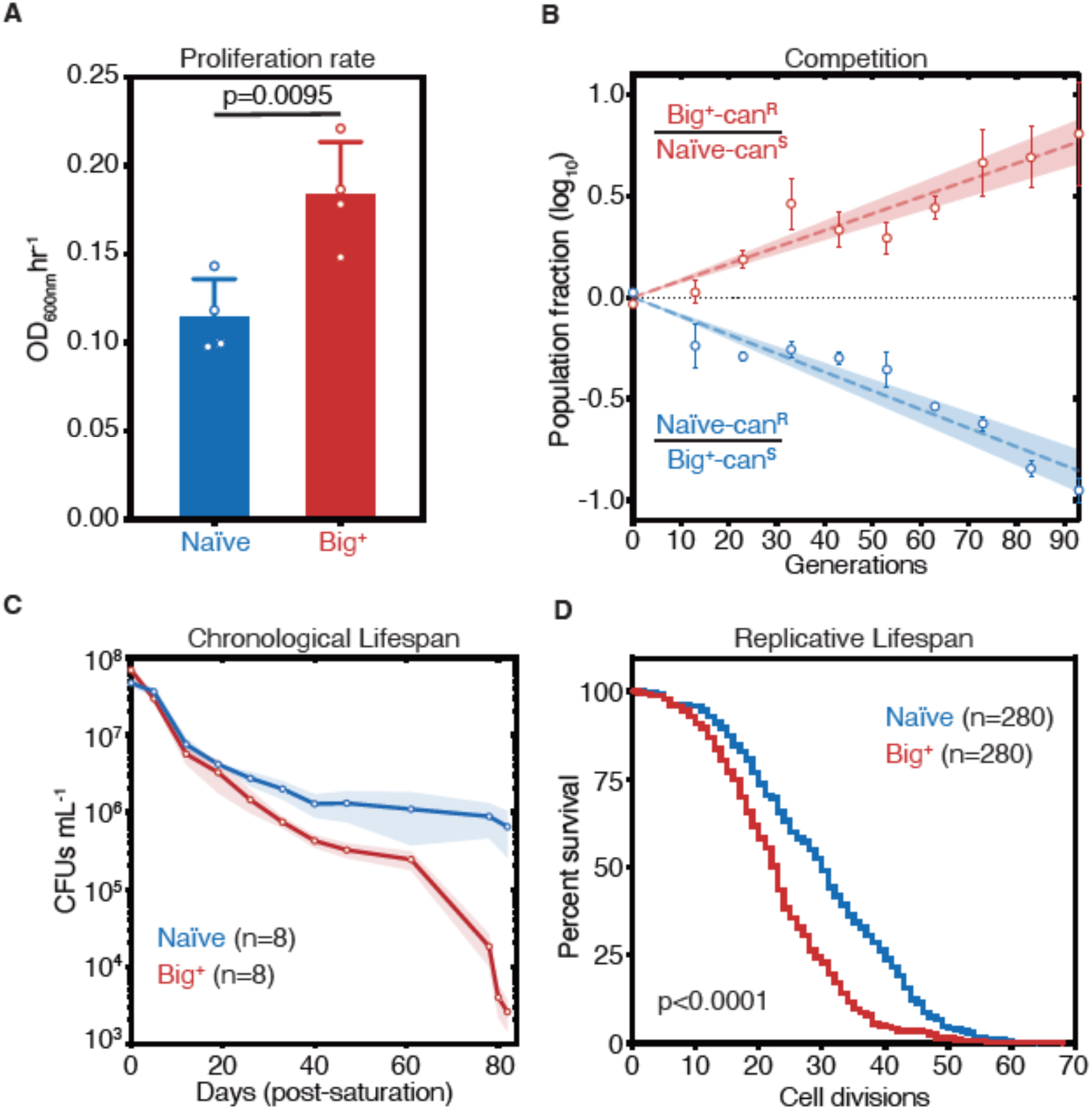
Cells bearing a prion-like epigenetic element live fast and die young. (**A**) Big^+^ cells proliferate faster than naïve cells. Bars represent mean of four replicates of maximum growth rate in YPD medium (measured by the peak of the derivative of the growth data), error bars are standard deviation, p=0.0095, unpaired t-test. (**B**) In a direct growth competition, Big^+^ cells outcompete naïve cells. Raw data with standard error bars; trend line is dashed line showing shaded standard error; four replicates were performed for each competition. (**C**) Big^+^ cells have a reduced chronological lifespan (CLS). Cells were grown to saturation in rich medium, and then transferred to nutrient poor medium and allowed to age for up to 80 days. Periodically samples were re-plated onto rich medium to measure remaining viability via colony forming units (CFUs). Thin lines are the average value from eight biological replicates with standard error represented by shading. (**D**) Big^+^ cells have a reduced replicative lifespan (RLS). Starting with virgin mother cells, at each cell division daughter cells were separated and the total number yielded was counted for each replicate. n=280 per strain, combined from three independent experiments. P value < 0.0001, by Gehan-Breslow-Wilcoxon Test. Median survival: naïve=30 generations, Big^+^=23 generations.

We next tested whether this growth advantage from the Big^+^ epigenetic element would be more pronounced during direct competition and oscillating nutrient availability—as a single-celled organism such as yeast often face in nature. To do so, we performed a competition experiment that encompassed periods of abundant nutrient availability followed by starvation. Using resistance to canavanine as a marker for naïve and Big^+^ strains, we mixed equal numbers of cells from each strain and propagated the mixed culture for close to 100 generations, diluting into fresh medium and measuring the fraction of the population that harbored the canavanine resistance marker every ∼10 generations (**Supplementary Fig. 1A**). In these experiments, cells harboring Big^+^ invariably outcompeted the genetically identical naïve cells that lacked it—they *live fast*—with a selection coefficient of nearly 1% (**Fig. 1B**). Competitions using reciprocally marked strains produced equivalent results. As a frame of reference, the fitness advantages that we measured for Big^+^ are larger than those conferred by >30% of non-essential genes (Breslow et al., 2008) and the vast majority of natural genetic variants that have been quantified (Jakobson and Jarosz, 2019; Jakobson et al., 2019; Sharon et al., 2018; She and Jarosz, 2018).

Given the close link between proliferation and aging in many organisms (Bitto et al., 2015), we investigated whether Big^+^ also influenced lifespan. Studies of aging in budding yeast have led to the discovery of numerous genes with conserved roles in aging of metazoans (Kaeberlein, 2010). These studies have measured two types of aging, both of which we tested here—chronological lifespan and replicative lifespan (Longo et al., 2012). Chronological lifespan is a measure of post-mitotic viability, during which cells cease division under starvation conditions, until nutrients become available again. These conditions occur commonly in the natural ecology of this organism (Landry et al., 2006). To investigate, we aged cultures of naïve cells and genetically identical cells harboring Big^+^ (**Supplementary Fig. 1B**). Over the course of 80 days, Big^+^ cells had progressively lower viability than matched, isogenic, naïve controls (**Fig. 1C**).

Replicative lifespan is a measurement of the number of cell divisions that yeast can undergo before death (**Supplementary Fig. 1B**). These experiments revealed a significant difference in replicative potential between Big^+^ and naïve cells—the median survival rate is reduced by seven generations (**Fig. 1D**, p<0.0001, Gehan-Breslow-Wilcoxon Test, and **Supplementary Fig. 1C**). This degree of lifespan shortening falls into the range seen for classic lifespan mutants: yeast lacking *SIR3* or *SIR4* have reductions of ∼3–4 generations, and yeast lacking *SIR2* by ∼10–11 generations (Kaeberlein et al., 2005; Kaeberlein et al., 1999). Thus, Big^+^ cells harboring this Pus4-induced element exhibit both a significantly decreased chronological lifespan and replicative lifespan: they *die young*.

### Modeling tradeoffs between proliferation and lifespan

We next investigated the adaptive value of the Big^+^ phenotype by quantitatively modeling its fitness consequences in fluctuating environments, where committing to a single strategy can impose limits on the long-term fitness of a population. For example, when nutrient-rich periods tend to greatly exceed starvation periods, the rapid growth of the Big^+^ phenotype might be favored, in spite of its die-young phenotype (**Fig. 2A**). But this same decision could be maladaptive if growth conditions skewed towards frequent and longer periods of nutrient scarcity—where cells that grow slower and die older would instead have an advantage (**Fig. 2B**).

**Figure 2.**
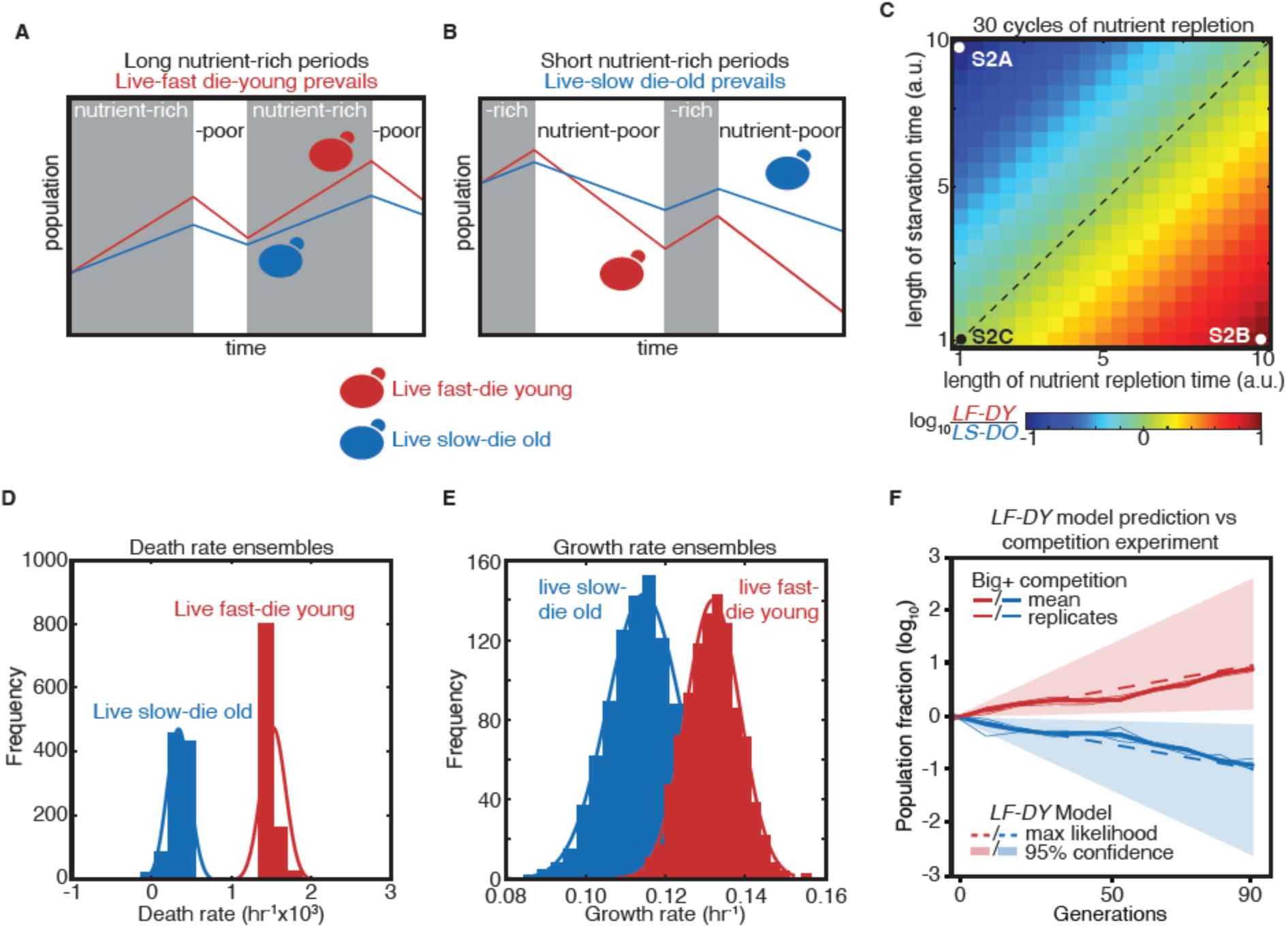
Modeling a reversible epigenetic live fast and die young strategy. (**A**) A live fast–die young epigenetic element is beneficial for survival in environments with regular, extended nutrient-rich periods. (**B**) A live slow–die old growth state is beneficial for survival during conditions of repeated and extended starvation. (**C**) Simulated final population fraction (ratio of *LF-DY* to *LS-DO*) after 30 cycles of nutrient repletion and starvation, assuming a 1% growth advantage, a 1% higher death rate, and equal starting population sizes. Note log scale. (**D**) Monte Carlo sampling of exponential decay constant and (**E**) exponential growth constant distributions used to generate the ensemble of simulations shown in, (**F**) Monte Carlo simulation (dashed lines) of growth competition between *LF-DY* to *LS-DO* cells under parameters sampled from experimental growth and lifespan measurements of the Big^+^ element (**Fig. 2D–E**). 95% confidence interval indicated by shaded areas. Shown in solid lines are the results of competitive growth between Big^+^ and naïve cells as shown in Fig. 1B (mean: bold line; n = 4 biological replicates: thin lines).

To quantify these tradeoffs, we considered a population in which individuals heritably adopted one of these two strategies (live-fast-die-young, *LF-DY*, like Big^+^ cells; or live-slow-die-old, *LS-DO*, like naïve cells), modeling their fates in alternating nutrient environments (**Fig. 2C**). These simulations suggested that *LS-DO* cells should outcompete *LF-DY* cells when the periods of starvation are much longer than periods of nutrient abundance (**Supplementary Fig. 2A**). By contrast, *LF-DY* cells come to dominate the simulated culture when periods of nutrient abundance are much longer than periods of starvation (**Supplementary Fig. 2B**). When periods of starvation and nutrient repletion are of equal duration, both populations are equally fit (**Supplementary Fig. 2C**). When we varied the growth advantage and lifespan cost of the *LF-DY* sub-population, and also the periods of nutrient availability and starvation, we noted that each phase space contained regimes in which *either* strategy could be advantageous (**Fig. 2C** and **Supplementary Fig. 2D**).

Regimes in which either the *LF-DY* or *LS-DO* strategies would be strongly adaptive (and the other maladaptive) arose frequently under physiologically relevant environmental parameters. Oscillations within these regimes (i.e. between feast and famine) are common in nature (Broach, 2012), and withstanding them is essential for survival. Theory predicts that reversible epigenetic mechanisms, such as prions, could confer a selective advantage in fluctuating environments (King and Masel, 2007). Importantly, our model demonstrates that this advantage could derive not only from a tradeoff between improved stress resistance and impaired growth in normal conditions, but also from a growth advantage in times of plenty coupled with a disadvantage under stress.

Notably, the growth advantages we observed in our competition experiment were quantitatively consistent with Monte Carlo simulations sampled from our experimental measurements of the death rates (from chronological lifespan measurements, **Fig. 2D**) and growth rates (from proliferation rate measurements, **Fig. 2E**) of individual cultures in nutrient starved and replete conditions, respectively (**Supplementary Fig. 2E**). That is, the adaptive advantages that we measured in competition were equivalent to those that we predicted for a hypothetical *LF-DY* population after dozens of generations (**Fig. 2F)**. Thus, selection on these properties could be alone sufficient to favor maintenance of the Big^+^ state under the conditions we examined.

### Big^+^ cells are large

Positive correlations between growth rate and cell size have long been noted (Johnston et al., 1977; Schaechter et al., 1958; Su and O’Farrell, 1998; Turner et al., 2012). To determine if the faster-growing Big^+^ cells were also larger, we examined them microscopically, employing a widely-used image masking pipeline (Carpenter et al., 2006). This allowed us to measure the sizes of thousands of cells from multiple biological replicates, defining size distributions for both naïve and Big^+^ populations.

During exponential growth, populations of Big^+^ cells had a similar size distribution to populations of naïve control cells. However, as the cultures became denser, naïve control cells remained the same size whereas isogenic haploid Big^+^ cells became larger (**Fig. 3A** and **Supplementary Fig. 3)**. To simplify these comparisons, we scored distributions by the fraction of very large cells that we observed (one standard deviation or larger than the mean naïve size; **Fig. 3B**, n=4,678 and 5,501 cells shown for naïve and Big^+^, respectively). This increase in mean area from 22.01 µm^2^ to 25.36 µm^2^ corresponds to a 23% larger volume (approximating the yeast cell as a sphere, naïve cell mean radius = 2.65 µm, Big^+^ cell mean radius = 2.84 µm). In Big^+^ cultures 33.1 ± (2.8)% of cells exceeded this threshold, whereas 16.5 ± (1.8)% did in naïve cultures (p=0.0011 by unpaired t-test; **Fig. 3C**).

**Figure 3.**
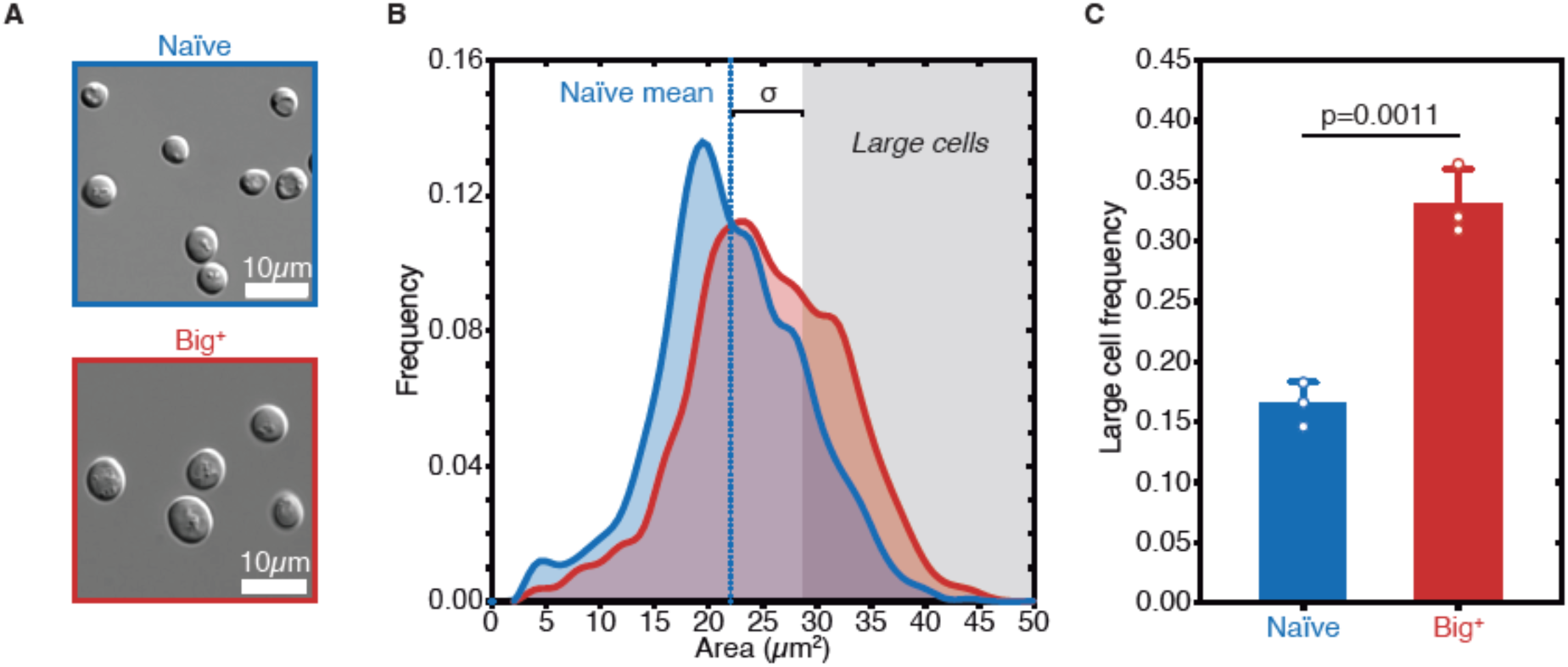
Big^+^ cells are large. (**A**) Micrographs of naïve and Big^+^ haploid yeast cells. (**B**) Cell size distributions for thousands of naïve and Big^+^ haploid cells (100% of distribution is shown, n=4,678 for naïve, n=5,501 for Big^+^, dotted lines indicate mean). Large cell threshold begins at one standard deviation above the naïve mean. (**C**) The frequency of haploid cells above the large cell threshold. Bars represent mean of three replicate strains, for which thousands of cells are measured for each strain, error bars are standard deviation. p=0.0011, unpaired t-test.

### [*BIG*^+^] is a prion transmitted through mating and meiosis

Big^+^ was originally induced by transient *PUS4* overexpression in a screen to identify prion-like epigenetic elements (Chakrabortee et al., 2016a). We therefore tested whether the increased cell size associated with this state was transmitted through genetic crosses with the unusual patterns of inheritance that characterize prion-based phenotypes (Brown and Lindquist, 2009; Cox, 1965; Wickner, 1994). We began by mating the large haploid Big^+^ cells to naïve haploids of the opposite mating type, selecting diploid cells, and measuring their size. The resulting diploids were significantly larger than those derived from control crosses with two naïve parents (**Fig. 4A–B**), establishing that the trait is dominant.

**Figure 4.**
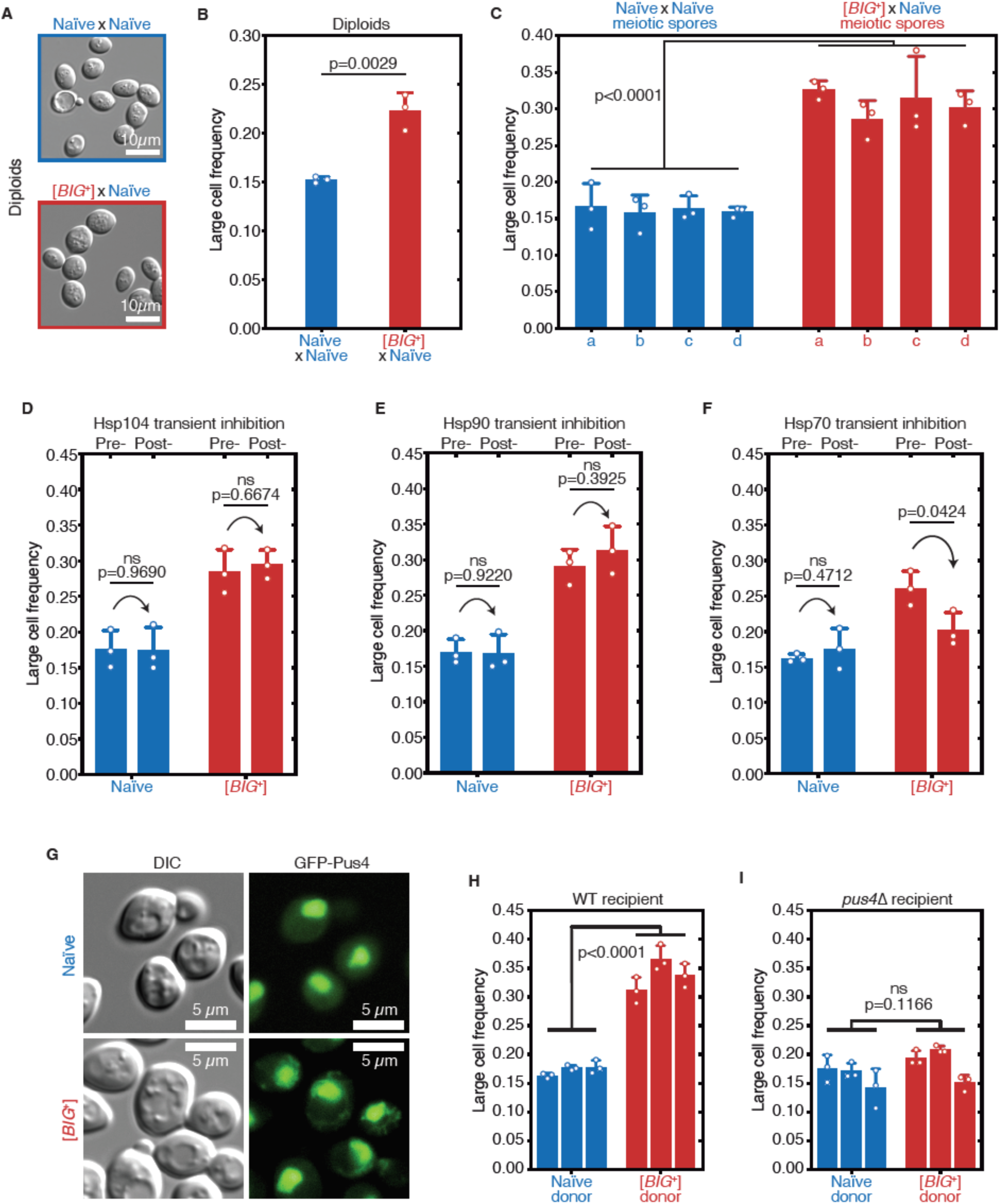
[*BIG*^+^] has prion-like patterns of inheritance. (**A**) Micrographs of diploid yeast cells resulting from crosses of naïve and naïve parents, or naïve and [*BIG*^+^] parents. (**B**) The frequency of diploid cells above the large cell threshold. Bars represent mean of three replicate strains, for which thousands of cells are measured for each strain, error bars are standard deviation. p=0.0029, unpaired t-test. (**C**) Inheritance of large cell frequency to all meiotic spores. Bars represent the mean frequency of cells above the large cell threshold from three replicates, for which thousands of cells were measured for each replicate, error bars are standard deviation. Difference between the means of four tetrad spores between naïve and [*BIG*^+^], p<0.0001, unpaired t-test. (**D**) Transient inhibition of Hsp104 chaperone activity using guanidinium hydrochloride (GdnHCl) does not heritably alter the cell size trait. Bars represent the mean frequency of cells above the large cell threshold from three replicates, for which thousands of cells were measured for each replicate, error bars are standard deviation. Control samples (left bars of each pair) were propagated in parallel on nutrient-matched agar plates not containing GdnHCl. Post-inhibition represents strains subjected to GdnHCl treatment followed by recovery prior to cell size measurements (**Materials and Methods**). Naïve p=0.9690, [*BIG*^+^] p=0.6674; unpaired t-test for both. (**E**) Transient inhibition of Hsp90 chaperone activity using Radicicol does not heritably alter the cell size trait. Bars represent the mean frequency of cells above the large cell threshold from three replicates, for which thousands of cells were measured for each replicate, error bars are standard deviation. Control samples (left bars of each pair) were propagated in parallel on nutrient-matched agar plates not containing Radicicol. Post-inhibition represents strains subjected to Radicicol treatment followed by recovery prior to cell size measurements (**Materials and Methods**). Naïve p=0.9220, [*BIG*^+^] p=0.3925; unpaired t-test for both. (**F**) Transient inhibition of Hsp70 chaperone activity by expression of a dominant negative allele of *SSA1* permanently eliminates the [*BIG*^+^] cell size trait. Bars represent the mean frequency of cells above the large cell threshold from three replicates, for which thousands of cells were measured for each replicate, error bars are standard deviation. Control samples (left bars of each pair) did not contain the *SSA1*^K69M^ constituitive expression plasmid but were propagated in parallel on non-dropout but otherwise nutrient-matched agar plates. Post-inhibition represents strains subjected to plasmid expression followed by plasmid removal and recovery prior to cell size measurements (**Materials and Methods**). Naïve p=0.4712, [*BIG*^+^] p=0.0424; unpaired t-test for both. (**G**) The expression pattern of Pus4 is altered in [*BIG*^+^] cells. (**H**) [*BIG*^+^] can be transmitted via cytoduction into a wild-type recipient cell, consistent with a prion-based mechanism. Each bar represents the mean frequency of cells above the large cell threshold from three biological replicates, for which thousands of cells were measured for each replicate, error bars are standard deviation. Bars for three independent cytoductants are shown for each donor strain. Difference between the means of the three cytoductants between naïve and [*BIG*^+^] donors: p<0.0001, unpaired t-test. (**I**) [*BIG*^+^] is not transmitted via cytoduction into a *pus4Δ* recipient cell, indicating that prion transmission depends on continuous endogenous expression of Pus4. Each bar represents the mean frequency of cells above the large cell threshold from three biological replicates, for which thousands of cells were measured for each replicate, error bars are standard deviation. Bars for three independent cytoductants are shown for each donor strain. Difference between the means of the three cytoductants between naïve and [*BIG*^+^] donors: p=0.1166, unpaired t-test.

We next investigated the meiotic inheritance of the large cell size trait. Because they are not driven by changes in nucleic acid sequence, prions have unusual patterns of inheritance that defy Mendel’s laws. In addition to dominance in genetic crosses, prion-based traits can be passed to *all* progeny of meiosis, in contrast to DNA-based traits, which are inherited by half (**Supplementary Fig. 4A**)(Garcia and Jarosz, 2014; Li and Kowal, 2012; Liebman and Chernoff, 2012; Wickner, 2016). We first mated control naïve cells to naïve cells of the opposite mating type, sporulated the resulting diploids, and dissected their meiotic progeny. We then grew clonal cultures of these haploid meiotic progeny and examined their size distributions, which were indistinguishable from their haploid naïve parents (**Fig. 4C**). In contrast, all cultures derived from the meiotic progeny of Big^+^ × naïve crosses were large (**Fig. 4C** and **Supplementary Fig. 4B**), a non-Mendelian pattern of inheritance that differs strongly from the expected behavior for genetic mutants or chromatin-based epigenetic elements, but is consistent with a prion-based mechanism of transmission. We therefore term this state “[*BIG*^+^]” (with capital letters indicating dominance and brackets denoting its non-Mendelian pattern of segregation).

### [*BIG*^+^] propagation requires the Hsp70 chaperone

The inheritance of prion-based phenotypes, in contrast to those driven by genetic mutations, is strongly dependent upon the protein homeostasis network (Garcia and Jarosz, 2014; Harvey et al., 2018; Liebman and Chernoff, 2012; Shorter and Lindquist, 2005). As a consequence, *transient* inhibition of molecular chaperones can lead to *permanent* elimination of prion-based traits. We therefore examined whether the large size of [*BIG*^+^] cells also depended on protein chaperone activity (**Supplementary Fig. 4C**). Transient inhibition of the Hsp104 disaggregase, which regulates the inheritance of many amyloid prions (Chernoff et al., 1995; Eaglestone et al., 2000; Halfmann et al., 2012; Shorter and Lindquist, 2004), had no effect on cell size in either naïve or [*BIG*^+^] cells (**Fig. 4D**). Transient inhibition of the Hsp90 foldase, which regulates the transmission of a different subset of prions (Chakrabortee et al., 2016a), also had no impact on [*BIG*^+^] transmission (**Fig. 4E**). By contrast, transient inhibition of Hsp70, via expression of a dominant negative *SSA1*^K69M^ allele (Chakrabortee et al., 2016a; Jarosz et al., 2014; Lagaudriere-Gesbert et al., 2002), caused [*BIG*^+^] cells to permanently lose their large size phenotype (**Fig. 4F**). Thus like other prions (Brown and Lindquist, 2009; Chakrabortee et al., 2016a; Chakravarty et al., 2019), and unlike genetic mutations, propagation of [*BIG*^+^] is dependent on the activity of this ubiquitous molecular chaperone.

We note that Hsp70 expression drops dramatically as yeast reach saturation and begin to starve (Werner-Washburne et al., 1989; Werner-Washburne and Craig, 1989), and also decreases as cells age (Janssens et al., 2015). These are two scenarios in which [*BIG*^+^] is disadvantageous. Therefore, environmental conditions that favor growth, during which Hsp70 is abundant, also favor prion propagation. By contrast, conditions known to reduce Hsp70 expression and thereby increase prion elimination are also those in which prion loss would be favored.

### Pus4 protein has a different expression pattern in [*BIG*^+^] cells

Acquisition of [*PRION*^+^] states often impacts the localization of the proteins that encode them. To visualize Pus4 expression in naïve and [*BIG*^+^] cells, we employed a strain in which an N-terminal GFP tag was appended at the endogenous *PUS4* locus (Weill et al., 2018; Yofe et al., 2016). We did not observe large fluorescent foci typical of canonical amyloid prions (Alberti et al., 2009). We did, however, observe altered localization of Pus4. In naïve cells Pus4 consistently localized to the nucleolus, as has been previously reported (Huh et al., 2003). In [*BIG*^+^] cells the protein was also present in the nucleolus. However, we also observed substantial fluorescence in a fragmented network throughout the cytoplasm (**Fig. 4G)**, establishing that the distribution of Pus4 protein is altered in [*BIG*^+^] cells. Although a high-resolution structure awaits determination, our data are consistent with an altered physical state of Pus4 in [*BIG*^+^] cells.

### Endogenous Pus4 is required for propagation of [*BIG*^+^]

Although [*BIG*^+^] was induced by a transient increase in Pus4, and was stable after elimination of the inducing plasmid, we wanted to exclude the possibility that this prior overexpression event might have established a positive feedback loop leading to an enduring increase in Pus4 levels. We therefore constructed naïve and [*BIG*^+^] strains with a seamless N-terminal 3X-FLAG tag endogenously encoded at the *PUS4* locus. Using immunoblots to detect the FLAG epitope in naïve and [*BIG*^+^] cells, we observed equivalent Pus4 levels, indicating that the phenotypes we observed in [*BIG*^+^] cells were not simply due to increased expression of this tRNA-modifying enzyme (**Supplementary Fig. 4D**).

Prion proteins can be inherited through the cytoplasm and do not require exchange of genetic material for propagation. To test this, we performed a cytoduction, in which we mated [*BIG*^+^] cells to naïve recipient cells of the opposite mating type carrying the *kar1-Δ15* mutation. Upon mating, this mutation prevents nuclear fusion, permitting mixing of cytoplasm, but not nuclei, between donor and recipient cells (**Supplementary Fig. 4E, Materials and Methods**) (Vallen et al., 1992). The transfer of [*BIG*^+^] cytoplasm into naïve recipient cells resulted in the transfer of the [*BIG*^+^] cell size phenotype (**Fig. 4H**). However, this was only the case for wild-type recipients. Naïve recipients lacking *PUS4* did not acquire the [*BIG*^+^] cell size phenotype (**Fig. 4I)**.

It remained formally possible that a multi-protein prion state could be maintained by other cellular factors, even if prion-based phenotypes depended on Pus4. Therefore, we tested whether transient loss of *PUS4* was sufficient to permanently eliminate [*BIG*^+^]. We first deleted *PUS4* from [*BIG*^+^] and naïve cells. Upon *PUS4* deletion, the sizes of the mutants derived from naïve and [*BIG*^+^] parents were equivalent (**Supplementary Fig. 4F**). Together with our cytoduction data, this suggested that continuous production of Pus4 is required to maintain this trait. Furthermore, *PUS4* deletion did not increase the size of naïve cells, suggesting that [*BIG^+^*] does not inactivate Pus4, in contrast to many well characterized prions that phenocopy loss-of-function alleles of their underlying proteins (Byers and Jarosz, 2014). Finally, we restored the *PUS4* gene to its native locus in these same cells by homology-directed integration. Even after re-introduction of *PUS4*, the size distributions of both populations remained equivalent (**Supplementary Fig. 4F**). Thus, both the expression and the propagation of the [*BIG*^+^] phenotype require the continual presence of a *PUS4* gene product.

These various lines of evidence—transmission to all meiotic progeny, dependence on molecular chaperones, altered expression pattern, and requirement for continuous expression of the protein that initiated the epigenetic trait—lead us to propose that [*BIG*^+^] is a protein-based element of inheritance, a prion, formed by the Pus4 pseudouridine synthase.

### Pseudouridylation is maintained in [*BIG*^+^] cells

Because loss of *PUS4* did not phenocopy [*BIG*^+^], but did block propagation of the prion, we wondered whether the catalytic activity of Pus4 was maintained in [*BIG*^+^] cells. To measure pseudouridylation in naïve and [*BIG*^+^] cells, we employed a qPCR-based method that capitalizes on the enhanced susceptibility of pseudouridine to reacting with CMC (1-cyclohexyl-(2-morpholinoethyl)carbodiimide metho-p-toluene sulfonate; **Fig. 5A**)(Lei and Yi, 2017). When pseudouridines are “labeled” by CMC, and these RNAs are used as templates for replication by reverse transcriptase, the enzyme generates nucleotide deletions and other mutations at these sites that can be detected by differences in the melting temperature of the derived nucleic acid duplexes, compared to non-pseudouridylated or unlabeled controls. Using this approach, we examined pseudouridylation of an archetypical Pus4 substrate, tRNA_AGC_ (Ala). When a pseudouridine is present and CMC is added, the melting curve shifts relative to an unlabeled control (no CMC). We observed a similar leftward shift in melting curves for both naïve and [*BIG*^+^] cells. In contrast, negative control cells missing the Pus4 protein, and therefore not pseudouridylated at U55, did not produce this shift (**Fig. 5B**). These data establish that Pus4-dependent modification of tRNAs is maintained in [*BIG*^+^] cells.

**Figure 5.**
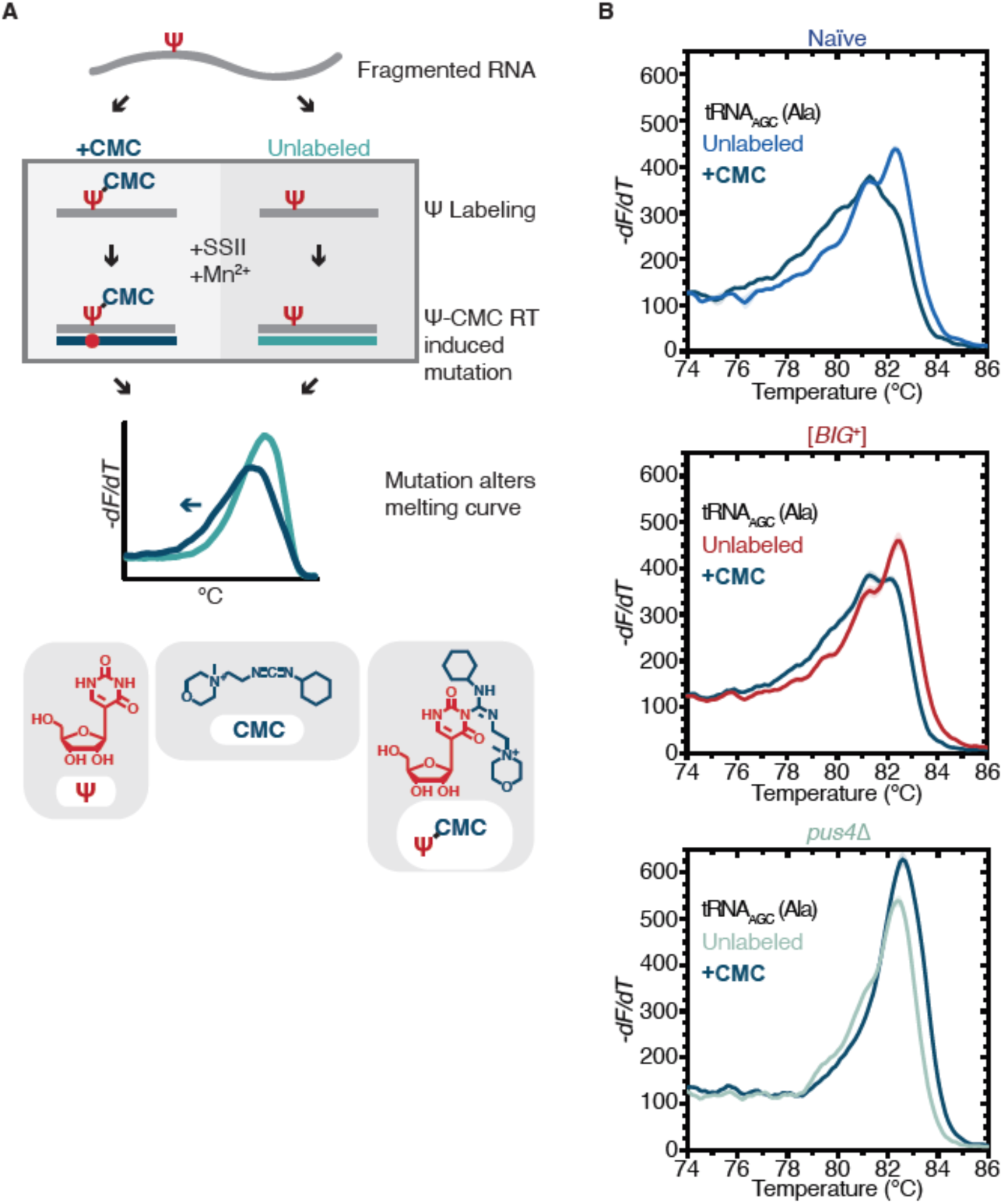
Pus4 activity is maintained in [*BIG*^+^]. (**A**) Radiolabeling-free, qPCR-based method for locus-specific pseudouridine detection. Figure adapted from reference (Lei and Yi, 2017). (**B**) High resolution melting curve analysis demonstrates that Pus4-dependent pseudouridylation of tRNA_AGC_ (Ala) is maintained in [*BIG*^+^] cells but not in cells that do not contain Pus4p. Top panel: naïve samples CMC-labeled (black) or unlabeled (blue). Middle panel: [*BIG*^+^] samples CMC-labeled (black) or unlabeled (red). Bottom panel: *pus4Δ* samples CMC-labeled (black) or unlabeled (torquoise). Solid lines representing melting curves are the mean of four replicates, with shaded areas representing standard error of the mean. The leftward shift of +CMC curves in naïve and [*BIG*^+^] but not *pus4Δ* samples indicate Pus4-dependent pseudouridylation of U55.

### Relative RNA levels are nearly unchanged in [*BIG*^+^] cells

In addition to its ubiquitous pseudouridylation activity on all tRNAs, Pus4 also modifies some mRNAs (Carlile et al., 2014; Lovejoy et al., 2014; Schwartz et al., 2014). Some have posited that pseudouridylation impacts mRNA stability (Zhao et al., 2017). To discern whether the phenotypes of [*BIG*^+^] cells might be due to relative changes in RNA levels, we performed RNA-sequencing. We grew naïve and [*BIG*^+^] cultures in YPD medium until late exponential phase, extracted total RNA, and depleted it of rRNA. Comparing five replicates each of naïve and [*BIG*^+^], the expression levels of only 15 genes changed significantly (thirteen decreased in expression and two increased, adjusted p-values < 0.1, Wald test, multiple testing correction with Benjamini Hochberg method), and these changes were modest (**Supplementary Table 2**). From this short list we did not observe any enrichments in gene ontology categories or pathway enrichments (YeastMine, *Saccharomyces* Genome Database (Cherry et al., 2012)). We conclude that the major effects of [*BIG*^+^] during exponential growth (e.g. growth rate and replicative lifespan) do not occur via major changes to steady state mRNA levels.

To investigate whether the relative abundance of tRNAs is perturbed in [*BIG*^+^] cells, we ran total RNA from naïve and [*BIG*^+^] cells (grown to late-exponential phase) on a nucleic acid fragment analyzer, and quantified tRNA levels relative to a similarly abundant RNA that is not a target of Pus4, 5.8S rRNA (158nt)(**Supplementary Fig. 5**). We observed no significant differences. While we cannot exclude other effects on tRNA function or the relative abundance of particular tRNAs, our data show that bulk tRNA abundance differences are likely also not responsible for the phenotypes of actively growing cells containing [*BIG*^+^].

### [*BIG*^+^] cells are resistant to inhibition of protein synthesis

Because the increased cell size and proliferation, and reduced lifespan of [*BIG*^+^] cells occur in the absence of major alterations to relative abundances of mRNA or tRNA, we wondered whether a change in protein synthesis might be responsible. To test this, we employed two inhibitors: 1) cycloheximide, an inhibitor of translational elongation (Baliga et al., 1969; McKeehan and Hardesty, 1969), and 2) rapamycin, a natural product macrolide that inhibits the TOR kinase, blocking a conserved signaling cascade that promotes protein synthesis (Beretta et al., 1996; Chung et al., 1992; Kuo et al., 1992; Price et al., 1992; Urban et al., 2007).

[*BIG*^+^] cells grew nearly two-fold better than naïve cells in a sub-inhibitory concentration of cyclohexamide that was sufficient to impair proliferation (0.05 µg/mL; 100 µg/mL is used in experiments that rapidly and completely arrest translation) (**Fig. 6A**; compare to growth rates in **Fig. 1A**). [*BIG*^+^] cells also proliferated faster in a concentration of rapamycin that inhibited growth (**Fig. 6B**), suggesting that the pathway may be more active in cells harboring the prion. This latter observation also provides a potential explanation for the decreased longevity of [*BIG*^+^] cells: loss of TOR pathway function is associated with extended lifespan in yeast (Dikicioglu et al., 2018; Fabrizio et al., 2001; Powers et al., 2006) and many other organisms including nematodes, fruit flies, and mice (Bjedov et al., 2010; Harrison et al., 2009; Robida-Stubbs et al., 2012).

**Figure 6.**
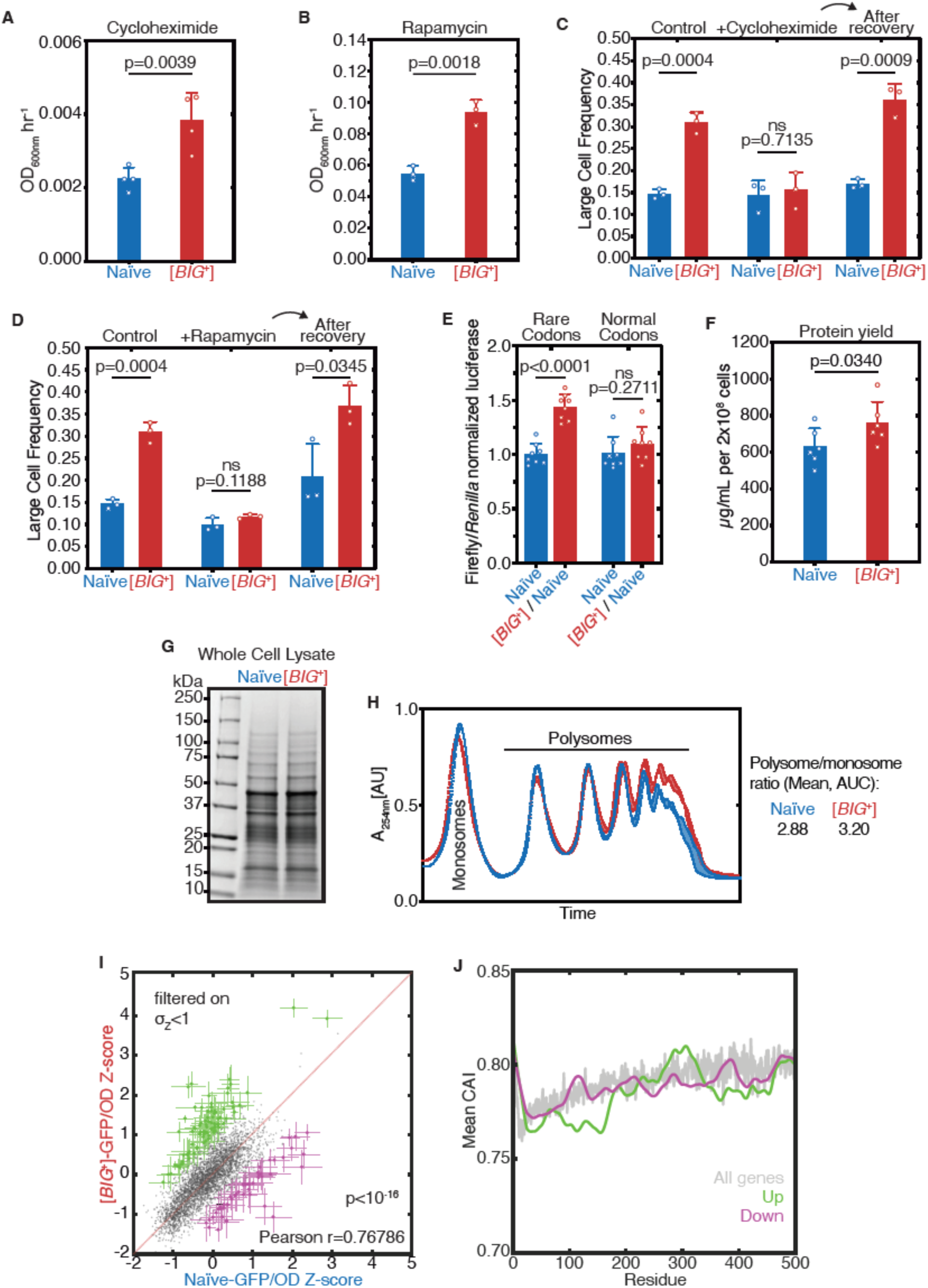
[*BIG*^+^] has altered protein synthesis. (**A**) [*BIG*^+^] cells are resistant to translation elongation inhibitor cycloheximide. Bars represent the mean of the maximum growth rate in YPD+cycloheximide (measured by the peak of the derivative of the growth data) of four replicates, error bars are standard deviation, p=0.0039, unpaired t-test. (**B**) [*BIG*^+^] cells are resistant to TOR inhibitor rapamycin. Bars represent the mean of the maximum growth rate in YPD+rapamycin (measured by the peak of the derivative of the growth data) of three replicates, error bars are standard deviation, p=0.0018, unpaired t-test. (**C**) [*BIG*^+^] cells grown in cycloheximide are not larger than naïve cells. However after recovery they regain this phenotype. After treatment, cells were subcultured in YPD for ∼75 generations before re-measuring the size in the absence of stress (see **Materials and Methods**). Bars represent the mean frequency of cells above the large cell threshold from three replicates, for which thousands of cells were measured for each replicate, error bars are standard deviation. Difference between the means of naïve and [*BIG*^+^]: YPD control p=0.0004; YPD+cycloheximide p=0.7135; YPD after recovery p=0.0009; unpaired t-test for all. (**D**) [*BIG*^+^] cells grown in rapamycin are not larger than naïve cells. However after recovery they regain this phenotype. After treatment, cells were subcultured in YPD for ∼75 generations before re-measuring the size in the absence of stress (see **Materials and Methods**). Bars represent the mean frequency of cells above the large cell threshold from three replicates, for which thousands of cells were measured for each replicate, error bars are standard deviation. Difference between the means of naïve and [*BIG*^+^]: YPD control p=0.0004 (same data presented in Fig. 6C as experiments were done in parallel); YPD+rapamycin p=0.1188; YPD after recovery p=0.0345; unpaired t-test for all. (**E**) [*BIG*^+^] meiotic progeny translate more of a Firefly luciferase reporter containing rare codons than naïve meiotic progeny do. This effect is not seen in an mRNA variant that encodes an identical protein but contains codons more frequently used in yeast. Bars represent mean normalized luciferase values (an invariable *Renilla* luciferase gene is co-expressed from the same plasmid) from eight replicates: rare codons p<0.0001, normal codons p=0.2711, unpaired t-tests for both. (**F**) [*BIG*^+^] cells produce more total protein per cell number than naïve cells, as measured by Bicinchoninic Acid (BCA) Assay. Bars represent the mean of six replicates, p=0.0340, unpaired t-test. (**G**) Coomassie stain of 15µg whole cell protein lysate from each strain suggests there are not major differences in the relative expression of the most abundant proteins in [*BIG*^+^] cells compared to naïve cells. (**H**) [*BIG*^+^] cells have more polysomes than naïve cells, as measured by polysome profiling. Lines for two technical replicates for each sample are shown, with the area between them shaded. Ratios (average of two technical replicates) were calculated by taking the lowest point between the monosome and disome peak as zero, and then calculating the ratios of the areas under the sum of the polysome peaks to that under the monosome peaks. (**I**) Plot showing proteome-wide GFP::protein fusion expression in [*BIG*^+^] cells compared to naïve cells, highlighting ∼130 proteins whose levels change. Each dot represents the mean of quadruplicate measurements of a single protein in naïve or [*BIG*^+^] cells: black dots are proteins that did not change significantly as measured by Z-score change of less than 1.0 (σ_Z_<1); green dots are protein fusions with higher fluorescence in [*BIG*^+^] cells; violet dots are protein fusions with lower fluorescence in [*BIG*^+^] cells. For colored dots, standard error of the mean is shown for both measurements from four biological replicates each. Pearson correlation of naïve and [*BIG*^+^] cells, r=0.76786, and p<10^-16^ indicates that most proteins have correlated expression levels. OD_600_ was adjusted based on known blank wells, and the GFP/OD600 measurements were normalized by Z-score ([*x*_i_-µ]/σ) within the naïve and [*BIG*^+^] populations independently. (**J**) Plot showing the protein residue number vs. the mean codon adaptation index (CAI) for all measured GFP-tagged proteins (grey line) and proteins whose levels were increased (green line) or decreased (violet line) in [*BIG*^+^] relative to naïve cells in the proteome-wide screen. Proteins whose levels were elevated in [*BIG*^+^] relative to naïve cells have a lower mean CAI in the 5′ ends of their mRNAs.

We next investigated whether these inhibitors impacted the size of [*BIG*^+^] cells. When grown in cycloheximide or rapamycin to saturation, [*BIG*^+^] cells were no longer large compared to naïve controls (**Fig. 6C–D**). Thus, unperturbed translation is necessary for the increased size of [*BIG*^+^] cells. These data could be explained by the inhibitors masking expression of the large cell trait, or by reversion of the [*BIG*^+^] prion. To distinguish between these possibilities, we sub-cultured cells that had been treated with cycloheximide or rapamycin and allowed them to recover in rich medium for several dozen generations. We then examined their size distributions. Cell populations derived from [*BIG*^+^] ancestors were once again significantly larger than naïve controls that we subjected to the same propagation regime (**Fig. 6C–D**). We conclude that [*BIG*^+^] depends on protein synthesis to augment cell size, but that the prion is stable to transient perturbations in translation.

### [*BIG*^+^] increases protein synthesis

Translation is rate-limiting for growth in nutrient-rich conditions (Kafri et al., 2016). We observed enhanced growth of [*BIG*^+^] cells in rich YPD medium (**Fig. 1A–B**). However, in synthetic defined medium with identical carbon source abundance (2% glucose), but fewer amino acids, nucleosides, and other nutrients for optimum growth (SD-CSM), [*BIG*^+^] cells did not grow faster than naïve controls (**Supplementary Fig. 6A**). These data, combined with the resistance of the prion cells to cycloheximide and rapamycin, suggested that protein synthesis might be enhanced in [*BIG*^+^] cells.

Because a major component of translational regulation is the efficiency with which each mRNA is translated by the ribosome, we examined the impact of [*BIG*^+^] on mRNAs encoded with different codons using luciferase reporter assays. We transformed [*BIG*^+^] cells and isogenic naïve control cells with dual-luciferase plasmids encoding both *Renilla* and firefly luciferase genes. We tested two versions of the firefly luciferase gene: the first contained the normal suite of firefly mRNA codons; the second produced an identical protein product, but via codons that are more rare in *S. cerevisiae*, reducing steady-state protein levels by ∼five-fold (Chu et al., 2014). Each plasmid also contained an internal control: the *Renilla* luciferase gene with its natural set of codons. The firefly reporter with “normal” codons did not produce more luciferase activity in [*BIG*^+^] cells than in naïve cells, when normalized to the *Renilla* control (**Fig. 6E**). However, the firefly reporter with rarer codons produced normalized luciferase levels approximately 50% higher in [*BIG*^+^] cells than in isogenic naïve control cells (**Fig. 6E**). These data suggest that [*BIG*^+^] cells may enhance translation of some proteins, especially those containing a greater frequency of rare codons. We observed these effects in both meiotic spores from naïve x [*BIG*^+^] crosses (**Fig. 6E**), and the original [*BIG*^+^] isolates (**Supplementary Fig. 6B).** Therefore the altered translation phenotype, like cell size, is inherited by all progeny of meiosis.

We next investigated whether [*BIG*^+^] had global effects on the proteome by isolating total protein from cells harboring the prion and naïve controls. We reproducibly obtained more total protein per cell from the [*BIG*^+^] cultures (**Fig. 6F**). We next loaded an equal mass of protein lysate from naïve and [*BIG*^+^] cells onto a denaturing polyacrylamide gel and separated them by electrophoresis. Coomassie staining of these gels showed no pronounced differences in banding patterns (**Fig. 6G**), suggesting that [*BIG*^+^] does not exert large changes on the composition of the major expressed portion of the proteome, proteins which are known to be efficiently translated (Gingold and Pilpel, 2011; Plotkin and Kudla, 2011).

We next performed polysome gradient analysis to assess global translation activity. We observed no change in monosomes or disomes in [*BIG*^+^] samples compared to naïve controls. However, polysomes—which are responsible for most protein synthesis (Noll, 2008; Warner and Knopf, 2002)—were increased in [*BIG*^+^] cells relative to naïve controls (**Fig. 6H**). In summary, we found that [*BIG*^+^] cells have higher levels of translation, which increases total protein output, and may particularly increase the levels of some proteins translated from mRNAs enriched with codons that are more rare in yeast.

### [*BIG*^+^] reduces time spent in G1 phase of the cell cycle

Conditions that enhance protein synthesis also tend to reduce the fraction of time that cells spend in the G1 stage of the cell cycle (Jorgensen and Tyers, 2004). This is due to the fact that commitment to S phase entry—budding of a daughter yeast cell—depends on sufficient production of proteins needed to replicate the genome and essential cellular structures. Accelerating the production of these factors can thus shorten this period. We measured the fraction of naïve and [*BIG*^+^] cells in the G1 phase of the cell cycle, by counting the fraction of unbudded cells (i.e. cells in G1). For naïve cells, 36.2% were in G1, whereas only 27.2% of [*BIG*^+^] cells were in G1, a ∼25% reduction (**Supplementary Fig. 6C**, p=0.0017, unpaired t-test). These data suggest that the cell cycle checkpoint for progression to S phase remains intact, and that a shortened G1 stage in [*BIG*^+^] cells is consistent with their increased protein synthesis. [*BIG*^+^] cells are not larger during exponential phase growth (**Supplementary Fig. 3**), suggesting that their cell size checkpoint remains intact. Cells in stationary phase, by contrast, which do not have the nutrient content needed to progress to S phase, may continue to accumulate mass at a faster rate, contributing to their larger size.

### [*BIG^+^*] cells enact an altered translational program

Because relative mRNA expression levels were only subtly altered in [*BIG*^+^] cells during exponential growth phase, but multiple measures of translation were increased, we considered the possibility that the phenotypes of the prion might be due to changes in protein levels of particular open reading frames. To test this idea, we capitalized on the dominance of [*BIG*^+^] in genetic crosses (**Fig. 4B**), mating haploid cells harboring the prion to a genome-wide collection of N-terminal seamless superfolder-GFP fusions (“SWAT” library; ∼5,500 ORFs; (Weill et al., 2018)). Equivalent matings between naïve strains and this genome-wide collection served as controls. To control for the larger size of the [*BIG^+^*] cells, we assessed protein levels in these diploid strains in terms of the relative GFP levels (normalized by OD_600_ and Z-scored) within the naïve and [*BIG^+^*] GFP-fusion collections separately. Mating and fluorescence measurements were performed in biological duplicate: the SWAT library was mated to two separate [*BIG*^+^] isolates alongside naïve controls. The reported OD_600_-normalized GFP levels are the mean of two technical duplicates of each biological duplicate. Protein levels measured in this way were generally well correlated between naïve and [*BIG^+^*] strains (Pearson’s *r* = 0.76; *p* < 10^-16^), in concordance with our results from electrophoresis of total cellular protein. Yet many proteins were up- or down-regulated in [*BIG^+^*] cells.

Of the 4,233 fusions whose abundance could be robustly quantified across the four replicates (σ_Z-score_ < 1 for both naïve and [*BIG^+^*]), ∼130 were differentially expressed in [*BIG*^+^] cells. Consistent with a bias toward enhanced translation, 81 were upregulated and 46 were downregulated (**Fig. 6I** and **Supplementary Table 3**). These proteins did not show any strong enrichment in physiochemical properties (see **Supplementary Text** for further discussion). Nor were they enriched in proteins encoded by the handful of known Pus4-pseudouridylated mRNAs (Carlile et al., 2014; Lovejoy et al., 2014; Schwartz et al., 2014). We also did not observe a widespread increase in expression of ribosomal proteins. We did, however, observe that proteins that were increased in [*BIG^+^*] cells had a modest decrease in their codon adaptation index (CAI) at the 5′ end of their mRNAs relative to all genes (**Fig. 6J**), indicating that cells harboring the prion might more efficiently translate these messages. The dip in CAI that is typical in the 5′ end of all genes, known as “translational ramping”, is thought to reflect the bias toward translational control near the beginning of ORFs (Frumkin et al., 2018; Tuller et al., 2010). (Lower CAI can both reduce the speed of elongation by requiring rarer tRNAs and lead to differences in mRNA structure that might also affect initiation or elongation efficiency.) These data are consistent with our luciferase reporter data in which rarer codons throughout the message output more protein in [*BIG^+^*] than in naïve cells (**Fig. 6E** and **Supplementary Fig. 6B**), as well as the resistance of prion cells to the elongation inhibitor cycloheximide (**Fig. 6A**).

Many hits were logically connected to the enhanced translation, increased size, and shortened lifespan of [*BIG*^+^] cells. The 81 upregulated proteins included multiple ORFs whose deletions are associated with decreased cell size (such as *PHO5, MRPS28, KAP122,* and *SWE1*) (Harvey and Kellogg, 2003; Jorgensen et al., 2002), increased chronological lifespan (*DIG2* and *UBX6*) (Garay et al., 2014), and extended replicative lifespan (*ENO1, MUB1, PHO87, PGM2,* and *YRO2*) (McCormick et al., 2015). Conversely, the 46 downregulated proteins included ORFs whose deletions are associated with increased cell size (*RNR4* and *RPL15B*) (Jorgensen et al., 2002; Perlstein et al., 2005) and decreased chronological lifespan (including *BUD23*, *SMI1*, *MHP1*, and *CLG1*) (Garay et al., 2014; Marek and Korona, 2013). Several proteins directly involved in translation control and ribosome biogenesis were also increased in [*BIG^+^*] cells, including *FHL1, MRPS28, MRPS16, RPL24B, RPS7A, and UTP10* (**Supplementary Fig. 6D**). The [*BIG^+^*]-regulated proteins also included 29 ORFs associated with differential sensitivity to rapamycin (including *SAS4, SNF1,* and *HCA4*) (Butcher et al., 2006; Dudley et al., 2005; Kapitzky et al., 2010), and 14 ORFs that are known genetic or physical interactors with TOR1 (including *GCD14* and *PAR32*) (Krogan et al., 2006; Varlakhanova et al., 2018). The functional breadth of these effects on the proteome, and their logical connection to factors involved in the control of proliferation, cell size, and lifespan suggest that the phenotypes of [*BIG^+^*] are likely derived from an altered translational regulation program that favors growth and proliferation at the expense of lifespan.

### Altered pseudouridylation in [*BIG^+^*] cells

Our finding that pseudouridylation on tRNA was maintained at similar levels in [*BIG*^+^] cells (**Fig. 5B**) led us to wonder whether mRNA substrates of Pus4 were similarly modified. The best-documented mRNA target of Pus4 is the translation elongation factor *TEF1/TEF2*—whose position U239 is robustly pseudouridylated in a Pus4-dependent manner (Carlile et al., 2014; Lovejoy et al., 2014; Schwartz et al., 2014). These paralogous genes encode identical copies of the EF-1 alpha translation elongation factor, which binds to aminoacylated tRNAs—specifically the T arm stem-loop that is pseudouridylated by Pus4 (Dreher et al., 1999)—and delivers them to the A-site of the ribosome during translation elongation (Schirmaier and Philippsen, 1984). Using the aforementioned qPCR-based method for detecting pseudouridylation, we also found that *TEF1/TEF2* was pseudouridylated in our wild-type cells (both naïve and [*BIG*^+^]), in a Pus4-dependent manner (**Fig. 7A**). Sanger sequencing verified that the modified position was identical to that previously annotated in the literature (**Supplementary Fig. 7A**).

**Figure 7.**
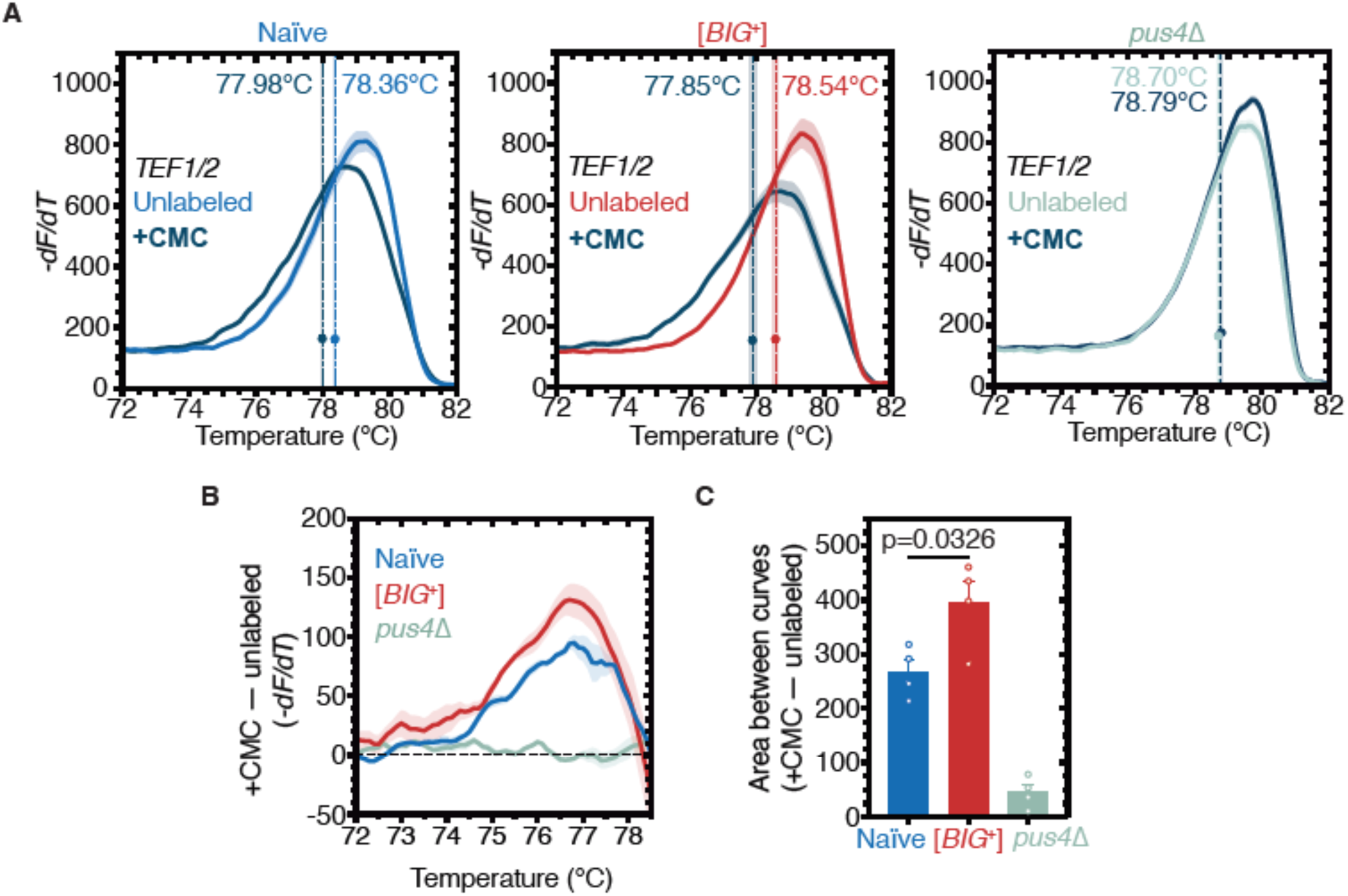
[*BIG*^+^] has elevated pseudouridylation. (**A**) High-resolution melting curve analysis shows that U239 is pseudouridylated in *TEF1/TEF2* mRNA in both naïve and [*BIG*^+^] cells but not in cells that do not contain Pus4p. Left panel: naïve samples CMC-labeled (black) or unlabeled (blue). Middle panel: [*BIG*^+^] samples CMC-labeled (black) or unlabeled (red). Right panel: *pus4Δ* samples CMC-labeled (black) or unlabeled (turquoise). Dots mark the geometric center of four replicates, bisected by a dashed line with shaded area representing standard error of the mean. The melting temperature (Tm) of this point is also displayed. Solid lines representing melting curves are the mean of four replicates, with shaded areas representing standard error of the mean. (**B**) The difference in melting temperature behavior (*df/dT*) between CMC-labeled and CMC-unlabeled *TEF1/TEF2* mRNA amplicons is larger in [*BIG*^+^] cells than in naïve cells, suggesting higher levels of pseudouridylation of U239 in [*BIG*^+^] cells. Solid line represents mean of four replicates, with shaded areas showing standard error of the mean. (**C**) The difference in area between the melting curves of CMC-labeled and CMC-unlabeled *TEF1/TEF2* mRNA amplicons is greater in [*BIG*^+^] cells than in naïve cells, suggesting higher levels of pseudouridylation of U239 in [*BIG*^+^] cells. Bars represent the mean of four replicates, error bars indicate standard deviation, p=0.0326, unpaired t-test.

The majority of studies on yeast prions have characterized them as decreasing or eliminating activity (Garcia and Jarosz, 2014). We recently discovered one notable exception, however, in which the [*SMAUG*^+^] prion can increase the activity of the protein that encodes it (Vts1; (Chakravarty et al., 2019). We thus examined if there was an altered level of pseudouridylaton of *TEF1/TEF2* mRNA—if altered levels affected protein activity, this could be one possible mechanism linked to the altered translation program we found in [*BIG*^+^] cells. By quantifying differences in the melt curve shift after CMC labeling—an analysis made simpler for *TEF1/TEF2* than for tRNAs by the absence of other pseudouridylated positions flanking U239—we observed an increase in the signal of pseudouridylation in [*BIG*^+^] cells relative to naïve (**Fig. 7B–C** and **Supplementary Fig. 7B**). Together these data demonstrate that the catalytic function of Pus4 is retained in [*BIG*^+^] cells, and can be enhanced for certain substrates, contrasting with the classical view of prions as being loss-of-function protein conformations. They also provide a novel example of how RNA modification can be epigenetically controlled.

### Epigenetic control of [*BIG^+^*]-like phenotypes in wild yeast

Finally we tested whether protein-based epigenetic control of cell size, protein synthesis, or localization is present in wild yeast populations. Protein chaperones are essential regulators of prion propagation in wild strains just as they are in laboratory strains (Halfmann et al., 2012; Jarosz et al., 2014). To block the passage of prions in wild yeasts, we transiently expressed a dominant negative variant (*SSA1*^K69M^) of Hsp70—the chaperone that is essential for [*BIG^+^*] propagation (**Fig. 4F**)—in twenty wild *S. cerevisiae* strains isolated from a variety of environments around the world (Cubillos et al., 2009; Itakura et al., 2019). We measured the size of cells before and after this chaperone curing and found four isolates—’273614N’, (clinical, Newcastle, UK); ‘L-1528’, (fermentation, Cauquenes, Chile); ‘DBVPG1373’, (soil, Netherlands); ‘BC187’, (barrel fermentation, Napa Valley, California)—that became smaller after curing (**Fig. 8A**).

**Figure 8.**
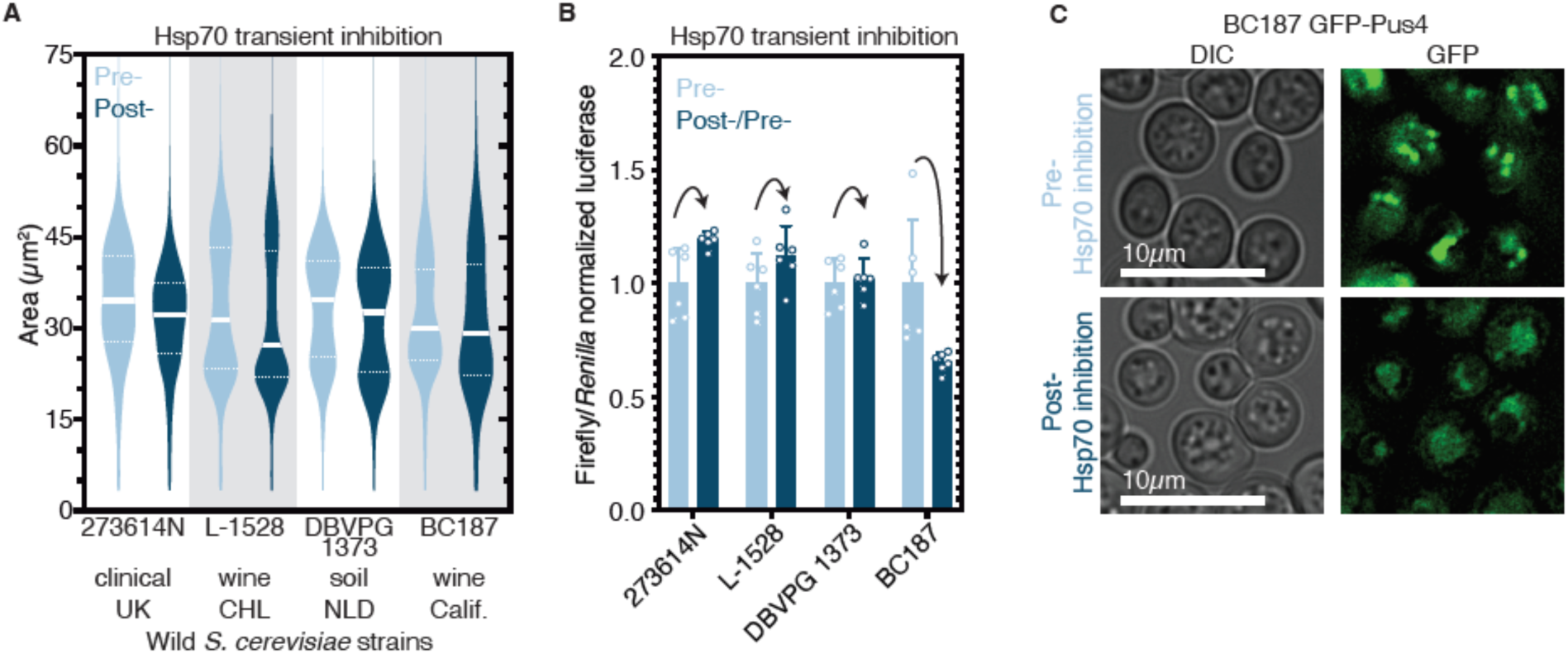
Epigenetic control of [*BIG^+^*]-like phenotypes in wild yeasts. (**A**) Transient inhibition of Hsp70 in diploid wild yeast strains from different niches around the globe leads to permanent reduction in cell size. Violin plots show all data from three biological replicates of each strain, light blue are cells before Hsp70 inhibition (“Pre-“), dark blue are cells after transient Hsp70 inhibition and recovery (“Post-“). Solid white line bisecting each distribution indicates mean; dotted lines indicate upper and lower quartiles. 273614N: clinical isolate from United Kingdom, Pre- vs. Post- p<0.0001. L-1528: wine isolate from Chile, Pre- vs. Post- p<0.0001. DBVPG 1373: soil isolate from the Netherlands, Pre- vs. Post- p<0.0001. BC187: wine isolate from California, Pre- vs. Post- p<0.0001. Kolmogorov-Smirnov test for all. (**B**) Transient inhibition of Hsp70 in wild yeast strains leads to permanent changes in protein synthesis capacity. Firefly reporter contains the “normal” suite of codons, which is normalized to internal control *Renilla* luciferase. Bars represent the mean of normalized luciferase values for six biological replicates, error bars are standard deviation. Light blue “Pre-“ are cells prior to Hsp70 inhibition, dark blue “Post-“ are cells after transient Hsp70 inhibition and recovery. A value above 1.0 for the dark blue bars indicates that the normalized translation capacity has increased after prion curing, a value below 1.0 indicates that capacity has decreased. 273614N p=0.0134, L-1528 p=0.1397, DBVPG 1373 p=0.7428, BC187 p=0.0125, unpaired t-test for all. (**C**) Transient inhibition of Hsp70 in BC187 wine isolate leads to permanent changes in Pus4p expression pattern, suggesting that its conformation may also be epigenetically regulated in wild strains. Prior to inhibition (top), cells show notable bright punctate structures; after transient inhibition and recovery (bottom), expression pattern becomes much more diffuse.

To test whether the four isolates that became smaller upon curing had altered protein synthesis, we transformed them with the same luciferase reporters we tested in laboratory [*BIG^+^*] cells. We normalized the firefly reporters with variable codons to the *Renilla* control for cured strains, and then normalized these to the corresponding uncured strain values. Two of the four isolates showed significant changes to translation (**Fig. 8B**). BC187 yielded the largest change upon curing, a ∼40% reduction in translation of a firefly containing a “normal” suite of codons, similar to what we had observed with [*BIG^+^*] cells generated in the laboratory.

We next examined the localization of Pus4 in BC187 and its Hsp70 ‘cured’ derivative. We transformed uncured and cured strains with a plasmid expressing GFP-tagged Pus4 protein, and imaged cells using epifluorescence microscopy. In uncured cells we observed distinct large puncta of Pus4. In contrast, in cured BC187 cells, the signal was far more diffuse; few cells had large puncta compared to uncured cells (**Fig. 8C**).

These observations suggest that epigenetic, and potentially prion-mediated control of mechanisms like the [*BIG*^+^] prion that we have described here, may be widespread in nature.

## DISCUSSION

Epigenetic inheritance is most commonly thought to be driven by enzymes that modify chromatin and DNA. Here we show that an enzyme that catalyzes epigenetic modification of RNA can itself be controlled by an extrachromosomal epigenetic process: a self-templating protein conformation that persists over long biological timescales. This prion-based mechanism engages an altered translational program to favor a ‘*live fast, die young*’ strategy.

RNA modifications can facilitate multiple steps of protein synthesis, including tRNA charging, ribosome biogenesis, and decoding (Sarin and Leidel, 2014). The epigenetic conformational control of an ancient RNA modifying enzyme that we have discovered provides a new translational control mechanism that strongly impacts growth, proliferation, and lifespan. These data further define a type of ‘recursive’ epigenetics, in which epigenetically transmissible information occurs via a protein that is itself an epigenetic regulator—a protein that chemically modifies RNA. We found that prion cells not only maintain pseudouridylation activity, but that activity can be increased, even without a detectable increase in *PUS4* expression.

Although the protein that drives [*BIG^+^*] modifies RNA, changes to relative RNA abundance do not appear to drive these growth or aging phenotypes. No major changes to relative tRNA or mRNA levels in actively growth cells are associated with [*BIG^+^*]. Instead, changes to the translational control of numerous genes are logically connected to these phenotypes.

It is difficult to predict a priori how the increased level of pseudouridylation in *TEF1/TEF2* mRNA might impact the protein’s activity. Due to a paucity of studies, the effects of pseudouridylation in ORFs is not well understood. Although the modification has been found to stabilize some mRNAs (Kariko et al., 2008; Nakamoto et al., 2017), recent evidence points to single sites slowing translation and altering decoding accuracy (Eyler et al., 2019). Given that our data are consistent with a change to translation elongation in [*BIG*^+^] cells, and the major non-tRNA substrate of Pus4 is the *TEF1/TEF2* mRNA, encoding a central elongation factor that binds to and escorts tRNAs to the ribosome, future studies should address whether Pus4-dependent modification of *TEF1/TEF2* mRNA plays a role in the phenotypes of [*BIG*^+^]. Apart from its pseudouridylation activity, the bacterial homolog of Pus4, *truB*, also harbors important tRNA chaperone activity (Keffer-Wilkes et al., 2016). It remains to be tested, however, if this activity is conserved in eukaryotic versions of the enzyme, and if so, what role if any that it plays in [*BIG*^+^].

Translation is a rate-limiting step for growth in many organisms (Polymenis and Schmidt, 1997; Sonenberg, 1993) and is often activated in human cancers (Sonenberg and Hinnebusch, 2009). Other prions in yeast also affect translation, including [*PSI*^+^] and [*MOD*^+^] (Baudin-Baillieu et al., 2014; Suzuki et al., 2012). However, in contrast to [*BIG*^+^], they lead to losses of their underlying protein activities, impairing translation and, in turn, growth in many conditions (Baudin-Baillieu et al., 2014; Cox, 1965; Suzuki et al., 2012). In contrast, in [*BIG*^+^] cells, translation is amplified. This is also notable for the fact that we have not found an example in the literature of a mutation that enhances translation under nutrient replete conditions. Moreover, [*BIG*^+^] did not require the amyloid severing activity of Hsp104 that is critical for propagation of [*MOD*^+^] and [*PSI*^+^], but rather the activity of a more generalist chaperone, Hsp70. A detailed description of the molecular conformation of Pus4 protein in [*BIG*^+^] cells, how it enables pseudouridylation activity, and how it promotes translation are ripe questions for future investigation.

Translation is also coupled to cell size, proliferation and lifespan (Ecker and Schaechter, 1963; Kaeberlein and Kennedy, 2007; Lloyd, 2013; Steffen and Dillin, 2016; Tanenbaum et al., 2015). Cell size, which is determined in large part by activity of the TOR pathway (Fingar et al., 2002; Zhang et al., 2000), has been inversely correlated with lifespan (Anzi et al., 2018; He et al., 2014; Yang et al., 2011), and older cells are larger (Egilmez et al., 1990). Moreover, molecules that extend lifespan, such as rapamycin, also influence cell size and/or proliferation by restricting the cell’s translation capacity (Beretta et al., 1996; Terada et al., 1994). Here we describe an epigenetic paradigm that links all of these fundamental cellular properties: translation, cell size, proliferation, and lifespan. In future studies we would like to explore to what extent the effect on lifespan may serve as material for selection to favor or disfavor [*BIG*^+^]. At present, we favor a hypothesis in which the aging defect of [*BIG*^+^] cells is due at least in part to pleiotropic consequences of their increased proliferation and translation capacity. Although these features have already been linked to aging in genetic studies, we also note that such theories of antagonistic pleiotropy in aging are not without controversy (Hughes and Reynolds, 2005).

Replicative lifespan of wild budding yeast strains has been measured and varies widely (Kaya et al., 2015). Variation has been associated with changes in oxidative phosphorylation, respiration, and differences in metabolite biosynthesis. Genetic screening has also offered some insight as to the genetic basis of this variability (McCormick et al., 2015). Lastly, genetic mapping efforts have identified polymorphisms underlying natural chronological lifespan variation (Kwan et al., 2011). The genetic architecture of natural lifespan, both chronological and replicative, remains obscure, however. Our data further demonstrate that it can be subject to strong epigenetic control.

We note that prior studies have characterized genetic links between cell size and lifespan in yeast—mutants that make cells larger tend to age faster, and older cells tend to be larger than younger cells (Neurohr et al., 2019; Yang et al., 2011; Zadrag-Tecza et al., 2009). [*BIG*^+^] provides an epigenetic mechanism to heritably alter these relationships. Exerting epigenetic, rather than genetic, control over basic cell growth behaviors could be valuable in the face of fluctuations between nutrient-replete and nutrient-poor conditions. Theory predicts that such mechanisms can have strong adaptive value when the frequency of fluctuations is rare relative to the generation time of the organism (King and Masel, 2007). In agreement with these inferences, our modeling quantitatively described the long-run selective advantage of this ‘*live fast, die young*’ prion state that we measured, illustrating the importance of considering not only steady-state phenotypes but also the ecological context in which prion states are expressed. Of particular relevance to this point is our data demonstrating that transient perturbation of Hsp70 activity can eliminate [*BIG*^+^]. Conditions in which the prion confers a growth benefit match those that promote its propagation. Conversely, conditions in which the prion is detrimental, shortening lifespan, are also those known to reduce chaperone expression, and could thereby cure the prion. The strong dependence of [*BIG*^+^] on chaperones therefore means that it is a natural epigenetic sensor of its environment.

Our data from wild yeast isolates demonstrates that cell size and protein synthesis are often under epigenetic control in nature. Exploring whether a [*BIG*^+^]-like epigenetic mechanism that promotes proliferation in nutrient replete conditions is conserved in metazoans is a major goal for the future. Indeed cancer cells also experience frequent oscillations in their environments: as tumors grow and metastasize, cells are exposed to shifting gradients of oxygen and glucose that influence their growth rate (Martinez-Outschoorn et al., 2017; Schito and Semenza, 2016). Regulation of cell physiology in these situations is best understood at the level of transcriptional changes due to the relative ease of profiling them, but multiple lines of evidence suggest that translational hyperactivation can also fuel pathological proliferation (Robichaud et al., 2019). Our discovery of a protein-based mechanism that changes cell growth via engagement of an altered translational program argues for greater investigation into the epigenetic control of post-transcriptional processes, in both normal biology and disease.

## ACKNOWLEDGEMENTS

We gratefully acknowledge Rebecca Freilich and Alan Itakura (Stanford University), Mike Harms (University of Oregon), and Gabriel Neurohr (ETH Zürich) for their review of the manuscript, and members of the Jarosz laboratory for helpful discussions. We thank Maya Schuldiner (Weizmann Institute) for the generous gift of the SWAT GFP library. We thank Kathrin Leppek and Maria Barna (Stanford University) for assistance with polysome gradients and profiling and the use of their instrument. We thank Rebecca Zabinsky and Thomas Lozanoski (Stanford University) for help with RNA-sequencing analysis. We thank Adam Fries for assistance with microscopy and the Genomics and Cell Characterization Core Facility for assistance with RNA quantification (Institute of Molecular Biology, University of Oregon). DMG was supported by postdoctoral fellowships from the NIH (F32-GM109680), Ford Foundation, and the Burroughs Wellcome Fund Postdoctoral Enrichment Program (award number 1015119), in addition to the University of Oregon, and a Pilot grant from the University of Washington Nathan Shock Center (NIH P30AG013280). EAC was supported by an NSF Graduate Research Fellowship. CMJ was supported by a postdoctoral fellowship from the NIH (F32-GM125162). This work was also supported by grants to DFJ from the National Institutes of Health (NIH-DP2-GM119140), the National Science Foundation (NSF-CAREER-MCB116762), a Searle Scholar Award (14-SSP-210), a Kimmel Scholar Award (SFK-15-154), and a Science and Engineering Fellowship from the David and Lucile Packard Foundation.

## AUTHOR CONTRIBUTIONS

DMG, EC, and DFJ conceived and designed the project. DMG, EC, CMJ, MT and AD performed the experiments and analyzed data. CMJ implemented the modeling. MK supervised the replicative aging experiments. DMG, EC, CMJ, and DFJ wrote the manuscript. DFJ and DMG supervised the project.

---

**Supplementary Figure 1.**
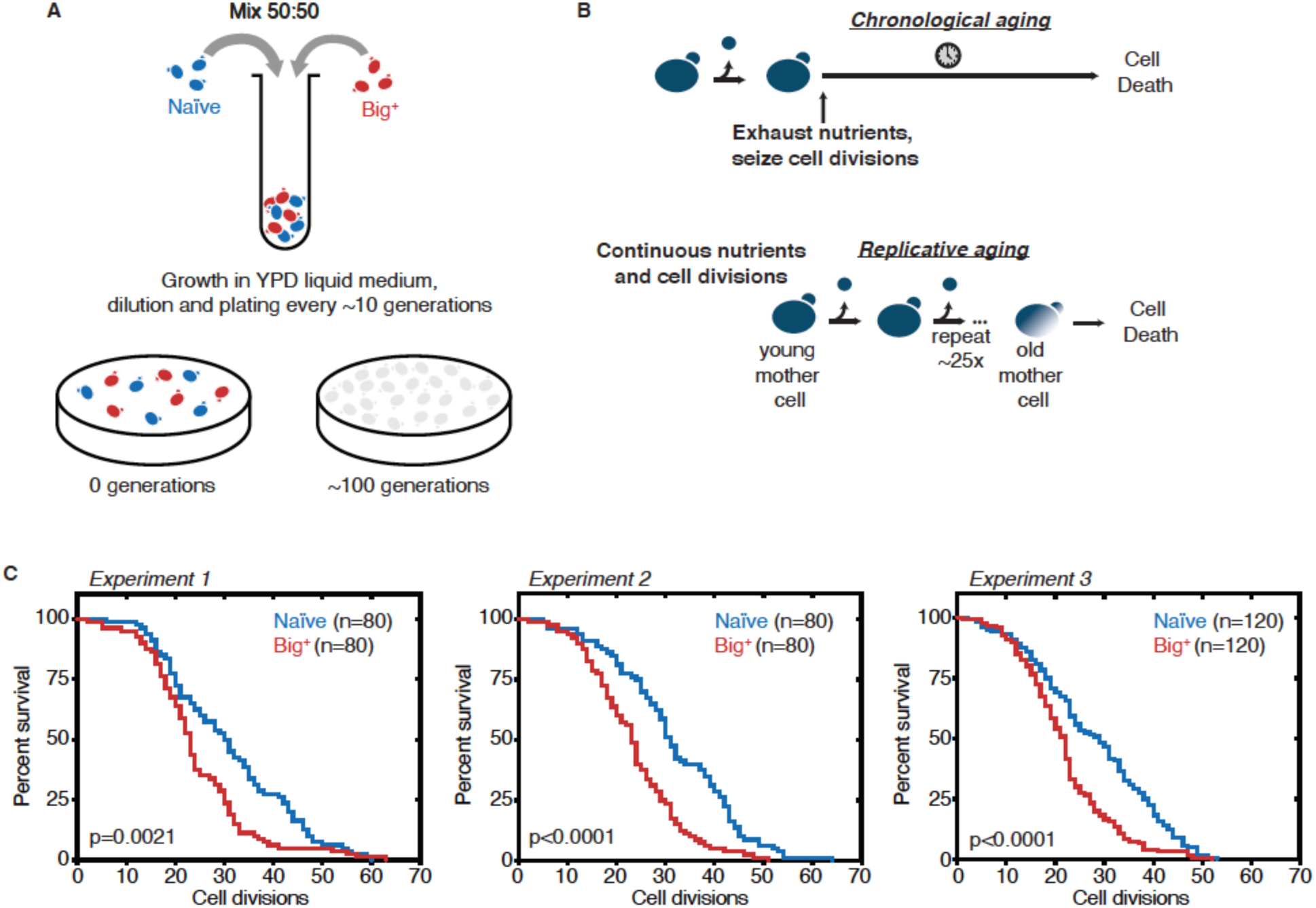
Probing lifespan of Big^+^ cells. (A) Experimental scheme for growth competition experiment associated with Figure 1B. (B) Experimental scheme for chronological and replicative lifespan measurements associated with Figure 1C–D. (**C**) Results from three independent RLS experiments, as combined into Figure 1D. Experiment 1: n=80 per strain, p value = 0.0021, by Gehan-Breslow-Wilcoxon Test. Median survival: naïve=30.5 generations, Big^+^=24 generations. Experiment 2: n=80 per strain, p value < 0.0001, by Gehan-Breslow-Wilcoxon Test. Median survival: naïve=31 generations, Big^+^=23 generations. Experiment 3: n=120 per strain, p value < 0.0001, by Gehan-Breslow-Wilcoxon Test. Median survival: naïve=29 generations, Big^+^=22 generations.

**Supplementary Figure 2.**
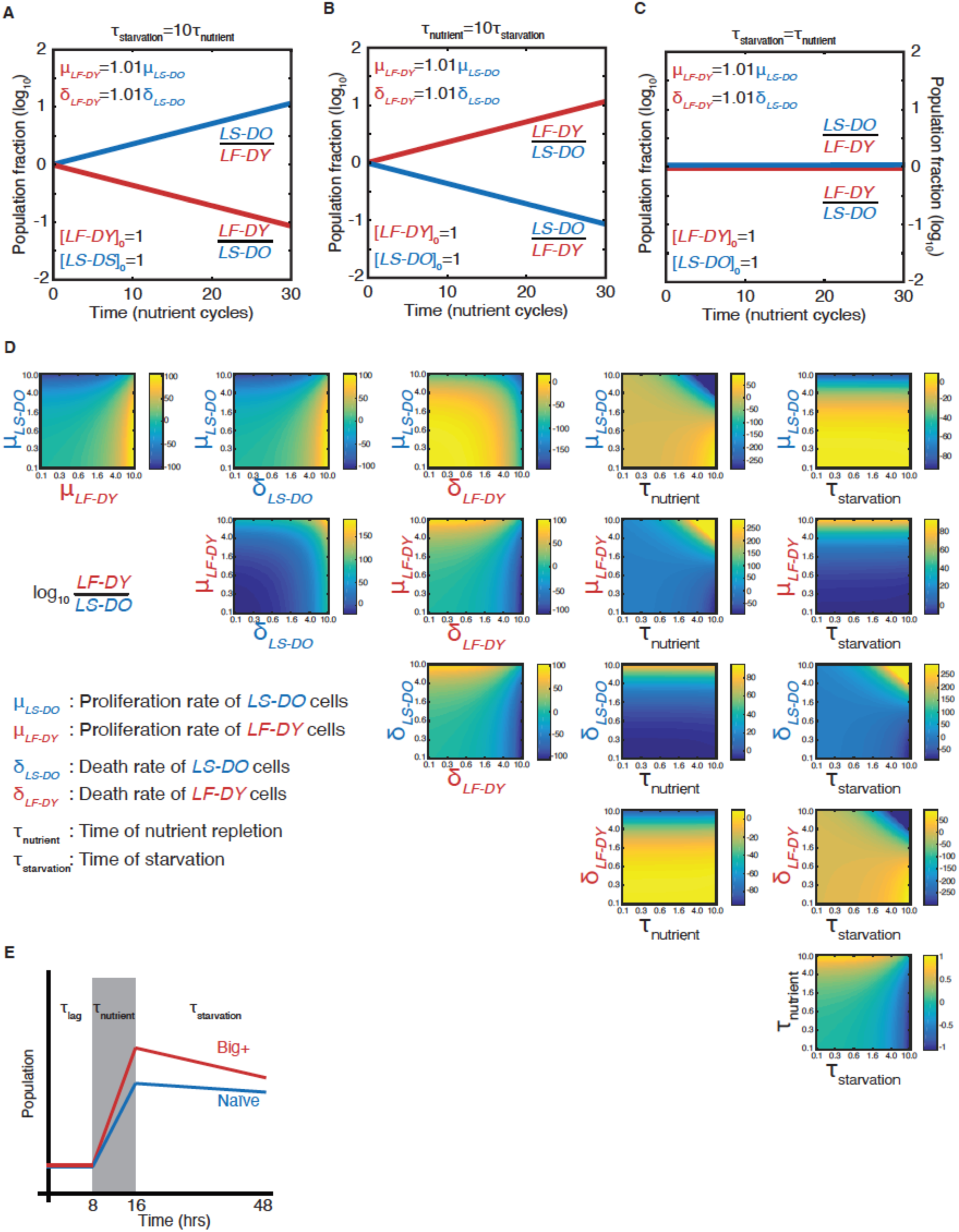
Modeling a reversible epigenetic live fast and die young strategy. Time-resolved simulations when (**A**) the period of starvation is ten times the period of nutrient repletion; (**B**) the period of nutrient repletion is ten times the period of starvation; and (**C**) the two nutrient regimes are of equal length. (**D**). Phase space representations of the simulated final population fraction (ratio of *LF-DY* to *LS-DO*) after 30 cycles of nutrient repletion and starvation, as in Fig. 2C. Indicated on the ordinate and abscissa of each panel are the parameters that were varied to generate each phase space. Parameters were varied over two orders of magnitude, as indicated. All other parameters were set to the baseline values as shown in **Supplementary Table 1**. (**E**) Schematic of parameters needed to model competitive growth experiment. During τ_lag_, we assume that there is no change in population ratio. During τ_nutrient_, we require the exponential growth constants, and during τ_starvation_, we require the exponential decay constants (assuming cell death is a first-order process).

**Supplementary Figure 3.**
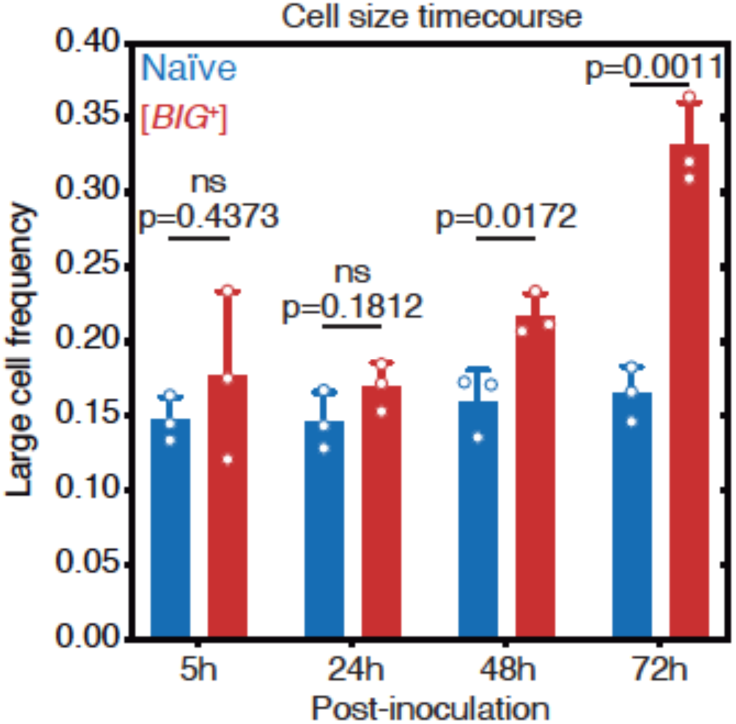
The large cell phenotype of Big^+^ emerges during the growth of a culture. The large phenotype of Big^+^ cells was stronger after three days of growth than after two days of growth (four days of growth yielded similar differences in cell size as for three days of growth, data not shown). The phenotype was not observed one day (24 hours) after inoculation or during the exponential growth phase (5 hours after inoculation). Bars represent the mean of three replicate strains—for which thousands of cells is measured for each—of the frequency of cells above the large cell threshold, error bars are standard deviation. Exponential growth (5 hours) p=0.4373, 24 hours p=0.1812, 48 hours p=0.0172, 72 hours p=0.0011; unpaired t-test for all.

**Supplementary Figure 4.**
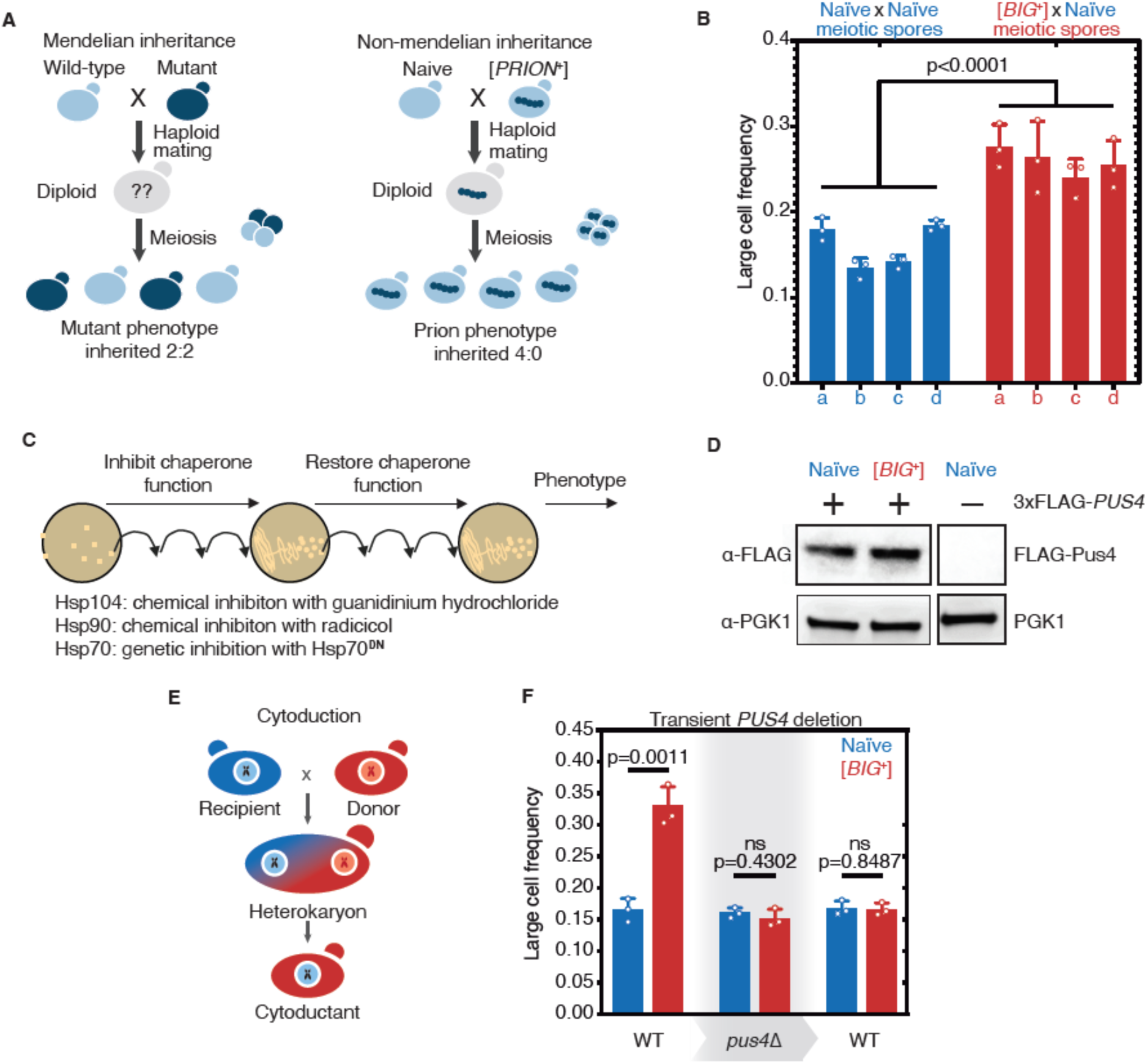
Tests for prion-like patterns of inheritance and dependence on Pus4 for [*BIG*^+^]. (**A**) In contrast to mutations, which when arising from one parent are inherited in half of the meiotic progeny, prion-based traits can be inherited in all meiotic progeny. (**B**) Additional pairs of tetrads are shown for naïve and [*BIG*^+^] crosses. Bars represent the mean frequency of cells above the large cell threshold from three replicates, for which thousands of cells were measured for each replicate, error bars are standard deviation. Difference between the means of four tetrad spores between naïve and [*BIG*^+^], p<0.0001, unpaired t-test. (**C**) Experimental scheme carried out to test the roles of three different protein chaperones in the propagation of [*BIG*^+^]. Cells were exposed to various chaperone inhibitors, then propagated without inhibition to allow cells to recover, and then tested for retention of the large cell phenotype that existed prior to inhibition. (See **Materials and Methods**.) (**D**) Pus4p is expressed at similar levels in naïve and [*BIG*^+^] cells. Naïve or [*BIG*^+^] haploid cells were crossed to a strain containing a seamlessly N-terminally 3xFLAG-tagged *PUS4* gene, and total protein lysate was harvested, of which 15µg was loaded onto a PAGE gel for each sample, and then probed with anti-FLAG, or anti-PGK1 loading control antibodies. Negative control lane (untagged strain) is from the same blot. (**E**) Prion-based traits can be passed through cytoduction that exchanges cytoplasmic material without exchange of nuclear material. (**F**) Transient deletion of *PUS4* blocks inheritance of the large cell trait from [*BIG*^+^] cells. Bars represent the mean frequency of cells above the large cell threshold from three replicates, for which thousands of cells were measured for each replicate, error bars are standard deviation. Differences between mean large cell frequencies of wild-type naïve and [*BIG*^+^] cells prior to deletion, p=0.0011; after deletion of *PUS4*, p=0.4302; after re-introduction of *PUS4*, p=0.8487; unpaired t-test for all.

**Supplementary Figure 5.**
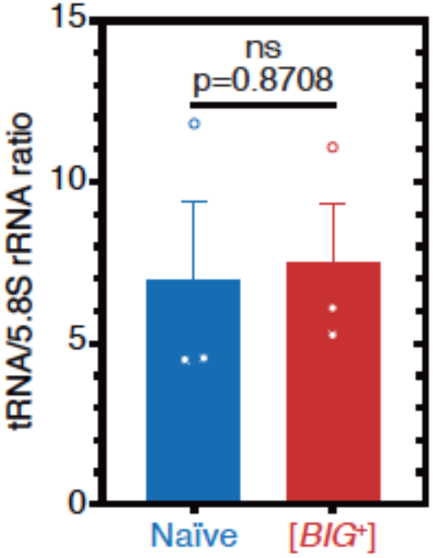
Unchanged tRNA levels in [*BIG*^+^]. tRNA levels are not different in [*BIG*^+^] cells compared to naïve cells when normalized to non-Pus4 target 5.8S rRNA (158nt). Measurements represent the mean of three replicates with standard deviation shown, p=0.8708.

**Supplementary Figure 6.**
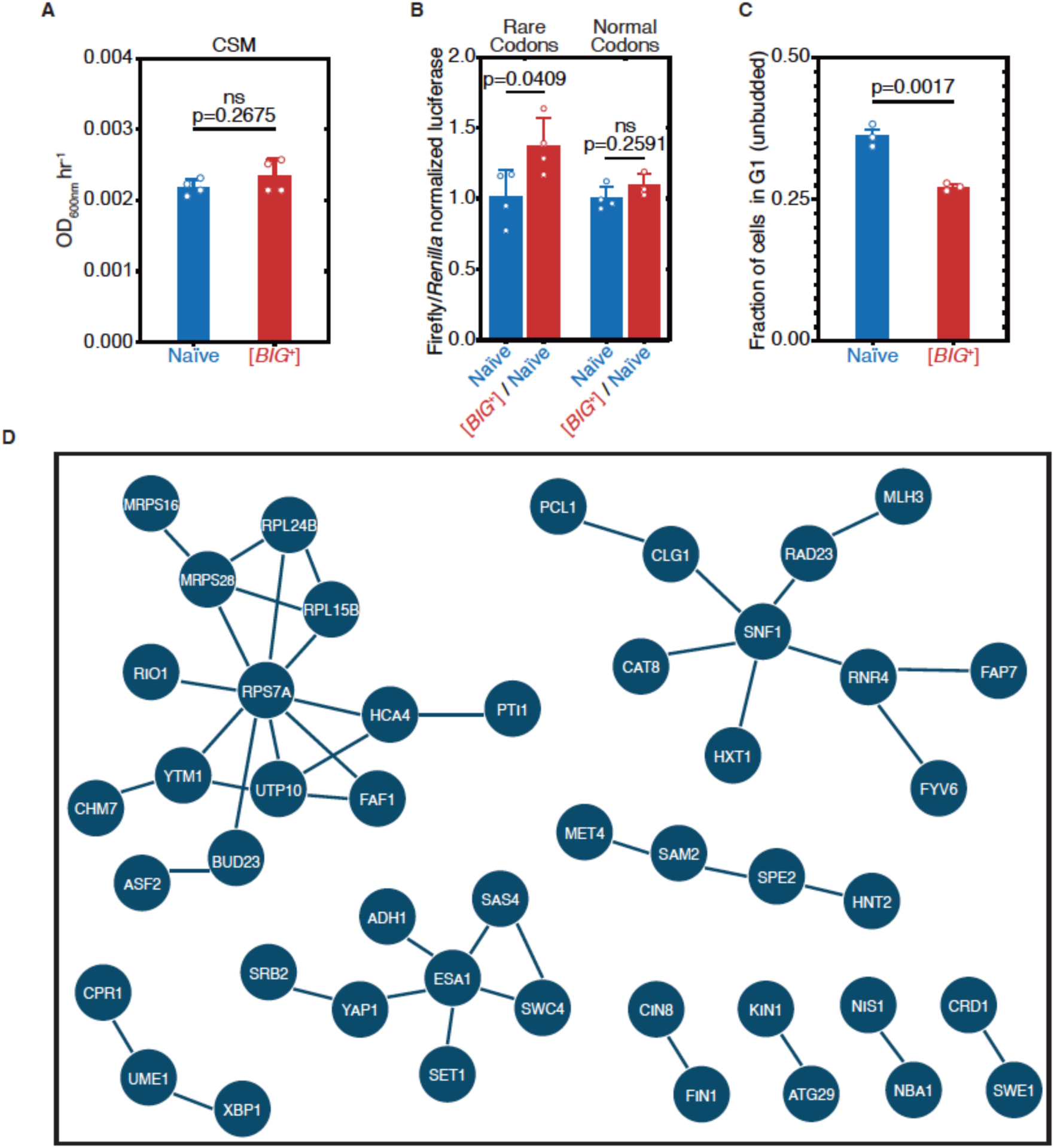
Altered cell cycle and translation in [*BIG*^+^]. (**A**) [*BIG*^+^] cells do not exhibit enhanced proliferation in SD-CSM, a less nutrient-rich medium than YPD (Fig. 1A). Bars represent mean of four replicates of maximum growth rate (measured by the peak of the derivative of the growth data), error bars are standard deviation, p=0.2675, unpaired t-test. (**B**) Original [*BIG*^+^] isolates translate more of a Firefly luciferase reporter containing rare codons than naïve cells do. This effect is not seen in an mRNA variant that encodes an identical protein but contains codons more frequently used in yeast. Bars represent mean normalized luciferase values (an invariable *Renilla* luciferase gene is co-expressed from the same plasmid) from four replicates: rare codons p=0.0409, normal codons p=0.2591, unpaired t-tests for both. (**C**) The ratio of unbudded cells (G1 phase of the cell cycle) to budded cells (G2 and S phases) is reduced in [*BIG*^+^]. Bars represent the mean of three replicates, error bars are standard deviation, p=0.0017, unpaired t-test. (**D**) Network representation of proteins whose levels change in [*BIG*^+^] vs naïve cells in GFP fusion screen (Fig. 6I and **Supplementary Table 3**), generated from STRING (string-db.org). Solid lines link proteins with genetic and/or physical interactions. The largest cluster—upper left— contains proteins involved in ribosome assembly and translation.

**Supplementary Figure 7.**
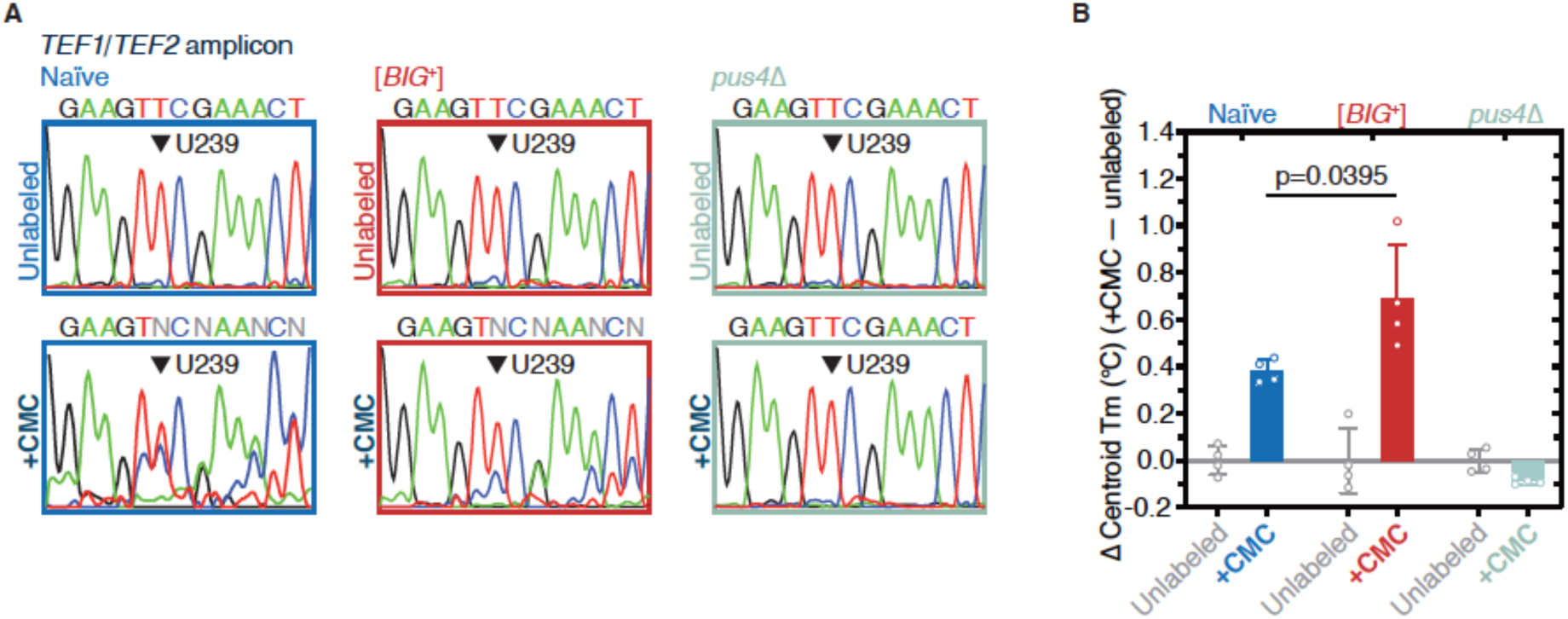
[*BIG*^+^] has elevated pseudouridylation. (**A**) Sanger sequencing profiles from control and CMC-labeled RNA from naïve, [*BIG*^+^], and *pus4*Δ cells. Naïve and [*BIG*^+^] samples show characteristic mixed nucleotide assignments at previously annotated pseudouridylated position U239 in *TEF1/TEF2* mRNA, as well as more variable assignments 3′ or this position, indicating the presence of a mixed population of amplicons containing CMC-pseudouridine induced mutations and deletions. (**B**) The difference in Tm between the curves of CMC-labeled and CMC- unlabeled *TEF1/TEF2* mRNA amplicons is greater in [*BIG*^+^] cells than in naïve cells, suggesting higher levels of pseudouridylation of U239 in [*BIG*^+^] cells. Each bar represents the mean of four technical replicates of the change in the centroid Tm, or the center of the distribution in both the x and y dimensions. Error bars are standard deviation. Difference between naïve and [*BIG*^+^], p=0.0395, unpaired t-test.

**Supplementary Table 1:**
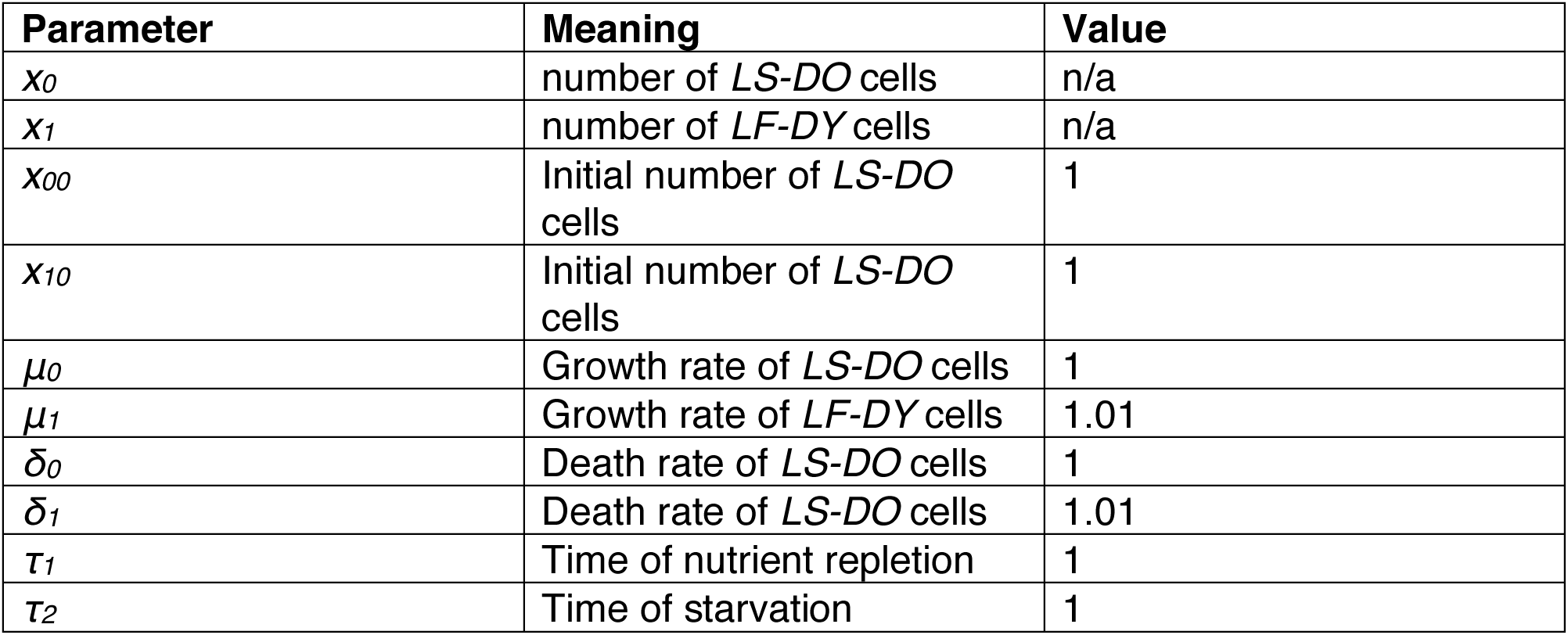
Parameter values used for the competitive fitness models shown in Figure 2 and Supplementary Figure 2.

| Parameter | Meaning | Value |
| --- | --- | --- |
| $x_0$ | number of *LS-DO* cells | n/a |
| $x_1$ | number of *LF-DY* cells | n/a |
| $x_{00}$ | Initial number of *LS-DO* cells | 1 |
| $x_{10}$ | Initial number of *LS-DO* cells | 1 |
| $\mu_0$ | Growth rate of *LS-DO* cells | 1 |
| $\mu_1$ | Growth rate of *LF-DY* cells | 1.01 |
| $\delta_0$ | Death rate of *LS-DO* cells | 1 |
| $\delta_1$ | Death rate of *LS-DO* cells | 1.01 |
| $\tau_1$ | Time of nutrient repletion | 1 |
| $\tau_2$ | Time of starvation | 1 |

**Supplementary Table 2: RNA-sequencing results**

See .xls file for TPMs for each gene in five replicates of naïve and five replicates of [*BIG*^+^]. Genes that showed significant differences are listed in a separate tab.

**Supplementary Table 3. Proteins whose expression is changed in [*BIG*^+^] cells.**

See .xls file

**Supplementary Table 4.** Yeast strains, plasmids and oligonucleotides used in this study. **See .xls file**

## Supplementary Text

We also examined the relationship between the proteins whose expression was altered in [*BIG*^+^] cells and other features related to gene expression, including mRNA secondary structure and pseudouridylation. Comparing the altered gene set to yeast transcripts with more double-stranded regions (Kertesz et al., 2010), no relationship emerged. We also compared the gene list to mRNAs that are pseudouridylated by Pus4 (Carlile et al., 2014; Lovejoy et al., 2014; Schwartz et al., 2014), but did not see any significant correlation or changes among the limited number targets that contained Pus4-dependent pseudouridylation sites in both studies, and which were present in the SWAT gene collection (thirteen genes total). We conclude that the changes we observe for the translation of ∼130 messages cannot be explained by Pus4-dependent pseudouridylation of their mRNAs. Given the limited reproducibility of mapping pseudouridylation sites transcriptome-wide in yeast from past studies, with large variability sample to sample (Safra et al., 2017; Zaringhalam and Papavasiliou, 2016), higher precision measurements in the future may offer the opportunity to compare anew those mRNAs pseudouridylated by Pus4 and their protein levels in [*BIG^+^*] vs naïve cells.

We also did not observe any relationship between genes with increased or decreased protein levels in [*BIG*^+^] and isoelectric point (pI), protein length, protein half-life, or GO category enrichments (process, component or function). We did however observe several properties that were altered in both the genes that went up and those that went down: hydropathy score (GRAVY)(Kyte and Doolittle, 1982), aromaticity, and codon adaptability index (CAI), codon bias, and frequency of optimal codons. We observed some amino acid enrichments as well, with genes going up in [*BIG^+^*] being depleted in leucine, phenylalanine, and proline, and those down depleted in phenylalanine and aspartic acid.

Finally, given that many of the effects that we measured were post-transcriptional, we also examined genes with altered GFP levels (significantly up or down) in [*BIG*^+^] cells using the MEME Suite, specifically for enrichment of predicted binding sites for RNA binding proteins (Bailey et al., 2009). We found that binding sites for the CCR4-NOT deadenylase complex were significantly enriched in our hits (motif alt ID CNOT4; consensus motif ACACAWA; adjusted p-value 0.0135).

## MATERIALS AND METHODS

### Model formulation

To model the fitness of an epigenetic element for which growth in nutrient-replete conditions is improved and survival in starvation conditions is worsened, we define the following growth equations in nutrient-replete conditions:

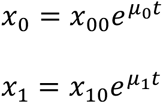

Where *x_0_* represents the population of naive cells and *x_1_* represents the population of [*BIG*^+^] cells. *μ_0_* and *μ_1_* are the growth rates of naive and [*BIG^+^*] cells, respectively. We neglect the lag and stationary phases of growth, as the ratio between populations does not change during this time.

Likewise, in starvation,

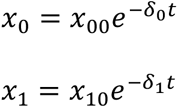

Where *δ_0_* and *δ_1_* are the death rates of naive and [*BIG^+^*] cells, respectively.

We can furthermore define the times of nutrient repletion and starvation as *τ_1_* and *τ_2_*, respectively. Thus, for each cycle of nutrient/starvation, we can define the following recursion relations:

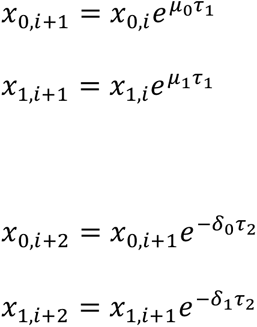

And we can write an analytical expression for the ratio of populations after one repletion/starvation cycle:

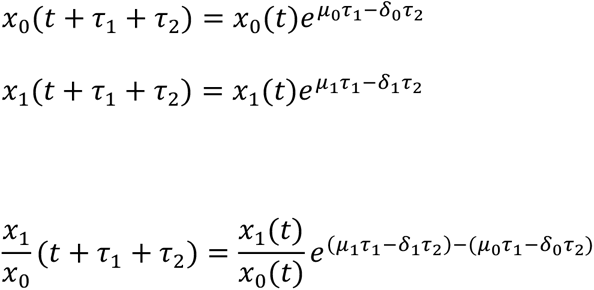

For the purposes of the model, we consider only the ratio between populations in the two epigenetic states (neglecting the total carrying capacity of the environment). The above defines a recursion relation from which we can predict the ratio of naive and [*BIG^+^*] cells after N cycles of nutrient repletion and starvation.

### Parameter estimation and prediction of competitive fitness

The free parameters defining the growth advantage and starvation disadvantage attributable to the epigenetic element in the model above (*μ_0_* and *μ_1_*; *δ_0_* and *δ_1_*) were determined by Monte Carlo sampling of independent measurements of the growth and death rates (**Fig. 2D–E**). To generate the ensemble of competitive fitness predictions shown in **Figure 2C**, we randomly sampled growth and death rates for naive and [*BIG^+^*] cells according to their experimentally determined distributions. The sampling was conducted 1000 times, and the prediction shown in **Figure 2F** is the median and 95% confidence interval of the resulting ensemble of model predictions.

### Code and data availability

All MATLAB code and data required to generate the model predictions is available at github.com/cjakobson/liveFastDieYoung. Code to generate the figures is available upon reasonable request to.

### Bacterial strain growth

Bacteria strains (for plasmid propagation) were cultured on LB agar or liquid (Research Products International (RPI), Mount Prospect, IL).

### Yeast strains

Yeast strains were cultured on either YPD agar or liquid (RPI) or SD-Ura (Sunrise Scientific, Knoxville, TN), unless otherwise indicated. Strains were stored as glycerol stocks (25% glycerol (Amersco, Solon, OH) in appropriate media) at −80 °C and revived on YPD or amino acid dropout media before testing. Yeast were grown in YPD at 30 °C on a TC-7 roller drum wheel (New Brunswick) unless indicated otherwise. Yeast transformations were performed with a standard lithium–acetate protocol (Gietz et al., 1992). The *pus4*Δ strain was sourced from the BY4741 MATa haploid knockout library (GE Dharmacon, Lafayette, CO).

### Strain constructions

Most diploids were constructed by crossing indicated BY4741 haploids to the BY4742 parental strain (ATCC, Manassas, VA) by mixing a bead of cells of each strain (from a single colony) together on a YPD plate and growing overnight at 30 °C. A small globule of this cell mixture was then re-streaked to single colonies on SD-Lys-Met agar plates to select for diploids.

Diploids constructed for the experiment presented in **Figure 4G** were made by crossing [*BIG*^+^] or naïve haploids to the seamless GFP-Pus4 haploid strain. Due to incompatibility of auxotrophic makers, diploids could not be selected on dropout plates from cell mixtures. Instead, a pool of the mated cells was grown in several successive competitions to allow diploids to outcompete haploid parents. Diploids were then isolated from single colonies and imaged as described below.

Sporulations were performed inoculating single diploid colonies in Pre-SPO liquid media (YPD with 6% glucose) for 2 days at room temperature on a roller drum wheel. Cells were then pelleted and washed twice in SPO media (1% Potassium Acetate (Sigma, St. Louis, MO), 320 mg CSM-Met powder (Sunrise Scientific), 20 mg Methionine (Sigma) per liter), and then diluted ten-fold into 3 mL cultures of SPO. These cultures were incubated on a rotary wheel for one week at room temperature, before dissecting tetrads on a Singer Instruments MSM400 (Somerset, England).

The 3XFLAG-Pus4 strains were constructed using PCR combined with the “Delitto Perfetto” method (Storici and Resnick, 2006). Canavanine mutants were constructed by pelleting 500*u*L of a saturated YPD liquid culture, resuspending it in 100 µLYPD and plating this on SD-Arg agar plates containing 60 µ*g*/mL canavanine (Sigma), and growing for 2 days at 30 °C. Single canavanine resistant colonies were picked and re-tested for resistance before further testing.

Cytoductions were performed as described in (Chakrabortee et al., 2016a). A BY4742 strain with a defective *KAR1* allele (*kar1-15*) was created as an initial recipient for cytoplasmic transfer. This allele prevents nuclear fusion during mating while permitting cytoplasmic transfer. The strain carries auxotrophic markers distinct from those in the putative [*BIG*^+^] or naïve donor strains, and was also converted to petite with growth on ethidium bromide (strains were grown in YPD with 25ug/mL ethidium bromide for ∼two-dozen generations before testing for growth on YP-glycerol). This allowed cytoplasmic transfer to be scored through the restoration of mitochondrial respiration, while selecting for auxotrophic markers unique to the recipient strain. The recipient and donor strains were mixed together on YPD-agar and grown overnight, followed by selection of heterokaryons and resulting haploid cytoductants on dropout media (selecting for the BY4742 recipient strain markers) containing glycerol as a carbon source. One more round of selection was used while replica-plating onto a dual-selection agar plate (SD-Lys-Met) to confirm that the colonies were not diploids. One additional round of propagation on a non-selective plate was performed before doing ‘‘reverse cytoductions,’’ which were performed in the same way except selecting for BY4741 auxotrophy in recipient naive strain. In the reverse cytoductions, the donors were naïve or putative-[*BIG*^+^] BY4742 *kar1-15* cytoductants from the first round, and the recipients were wild-type or *pus4Δ* naïve BY4741 petite cells.

Transient *PUS4* deletion experiment strains were made by the Delitto Perfetto method (Storici and Resnick, 2006). After deletion of *PUS4*, strains were propagated for ∼75 generations before phenotyping. The *PUS4* gene was re-introduced by homologous recombination.

### Growth assays

Biological replicates of each yeast strain were pre-grown in rich media (YPD). We then diluted these saturated cultures 1:20 in sterile water and then inoculated 3 µL into 96-well humidified plates (Nunc Edge Plates (Thermo Scientific, Waltham, MA)) with 150 µL of YPD or SD-CSM per well. Cycloheximide (Sigma) was added to growth media at 0.05µg/mL. Rapamycin (LC Laboratories, Woburn, MA) was added to growth media at 10µM. Cell growth was monitored with continuous measurements of OD_600_ (∼every 10 minutes) at 30°C over 96 hours using BioTek Eon or Synergy H1 microplate readers (Winooski, VT). Timepoints plotted in bar graphs correspond to the maximum proliferation rates calculated from growth data.

### Measurement of chronological lifespan

For each strain, four single colonies were picked from freshly streaked YPD plate, and grown in 5mL of Pre-sporulation media for 3 days on a roller drum wheel (New Brunswick Scientific, Edison, NJ) at 30°C. Cultures were then pelleted and washed once with SPO media, and resuspended in 5mL of SPO media, and placed back on the roller drum wheel at 30°C. On days indicated, a dilution was made of each replicate to achieve dozens to hundreds of colonies on a YPD plate, which were then counted using a colony counter (Synbiosos Acolyte, Frederick, MD). Dilution was ∼100,000X at early stages of experiment, and later on was sometimes empirically determined after significant cell death. We note that aging the cells in SPO did not lead to significant acidification (pH of old cultures was found to be >5), as has been reported for cells aged in YPD, which contains high levels of glucose that upon metabolism leads to secretion of organic acids (Murakami et al., 2011).

### Measurement of replicative lifespan

Replicative lifespan (RLS) was assessed using the standard method of isolating virgin cells on agar YPD (2% glucose) plates, and then separating their daughter cells at each cell division by micromanipulation and counting the total number of daughters produced by each mother cell (Steffen et al., 2009; Wasko et al., 2013). Strains were streaked from glycerol stocks onto YPD plates, and allowed to grow at 30°C until individual colonies could be selected for each strain. Colonies were lightly patched onto fresh YPD overnight and twenty cells were isolated by microdissection from each patch. These cells were incubated at 30°C for approximately two hours until they had formed daughter cells, at which time individual virgin daughter cells were selected and arrayed as previously described (Steffen et al., 2009) for lifespan analysis. From these, daughter cells were removed by microdissection and counted approximately every 2 hours during the day. Plates were maintained at 30°C during the day and placed at 4°C overnight. At least four independent replicates (arising from different colonies) of twenty individual mother cells each were measured for each strain.

### Curing

Three regimes of chaperone inhibition were tested: 1) transient exposure to a dominant negative version of Hsp70 (Ssa1) to inhibit its activity (Chakrabortee et al., 2016a; Jarosz et al., 2014) 2) transient exposure to Radicicol (LC Laboratories) to inhibit Hsp90 activity (Chakrabortee et al., 2016a) 3) transient exposure to Guanidinium Hydrochloride (Sigma) to inhibit Hsp104 activity (Ferreira et al., 2001).

Regime 1 was performed by transforming cells with a plasmid, PDJ169, harboring a dominant negative version of Hsp70 (Ssa1) as described previously (Chakrabortee et al., 2016a; Jarosz et al., 2014; Lagaudriere-Gesbert et al., 2002). Transformants were picked and re-streaked by hand or replica-pinned using a Singer HDA robot a total of twelve times on SD-Ura to promote Ssa1^DN^ expression. (Anecdotally, we note that prion phenotypes are frequently cured with fewer than twelve restreasks, but for reasons of technical throughput, twelve were used in this experiment.) Then plasmids were eliminated by plating on media containing 5-fluoroortic acid (SD-Ura + 0.1% 5-FOA + 50 µg/mL uracil) and plasmid loss was verified by replating on SD-Ura. Colonies were then tested for elimination of prion phenotypes. Tested strains were compared to control strains that were restreaked in parallel on SD-CSM plates.

Regime 2 was performed by replica-pinning cells six subsequent times on YPD agar plates containing 5 µM radicicol, using a Singer HDA robot. After re-platings, cells were plated back onto YPD two subsequent times to facilitate recovery before being tested for elimination of prion phenotypes. Tested strains were compared to control strains that were replica-pinned in parallel on YPD plates.

Regime 3 was performed like Regime 2 but with SD-CSM plates containing 0.5 g/L Guanidinium Hydrochloride. Tested strains were compared to control strains that were replica-pinned in parallel on SD-CSM plates.

Wild yeast strains were cured by transforming a uracil-selectable 2micron plasmid (PDJ1222) encoding the aforementioned dominant negative version of Hsp70 (Ssa1), under control of a constituitive promoter (*pGPD*). Transformants were passaged on selective media five times to allow growth of single colonies. Transformants were then passaged three times on non-selective media (YPD) to permit plasmid loss, which was confirmed by the lack of growth on selective media (SD-Ura).

### Strain competitions

Single colonies were used to inoculate 5 mL YPD cultures which were grown for 3 days on a roller drum wheel at 30°C. Cells were diluted 1000-fold and then mixed in equal volumes to form 50:50 mixtures of either of the following: Naïve Can^S^ and [*BIG*^+^] Can^R^; or Naïve Can^R^ and [*BIG*^+^] Can^S^. Before mixing cells, saturated cultures were measured to have near equal cell densities, and “time zero” measurement was made by plating the initial cell mixture and counting the number of Canavanine resistant colony forming units (CFUs) relative to total CFUs. These initial strain mixtures were then grown for 2 days on a roller drum wheel at 30°C, after which cells were diluted 50,000-fold or 25,000-fold and plated on YPD or canavanine plates, respectively. These plates were grown at 30°C for 2 days before counting colonies. The liquid cultures were diluted 1:1000 in 5 mL of fresh YPD, and this process was repeated nine more times. Swapping of canavanine resistance between naïve and [*BIG*^+^] was done to correct for the potential of canavanine resistance to influence cell growth, however in our experiments the differences were negligible. Numbers of canavanine-resistant and total colonies were compared relative to number of cells plated to determine the number of naïve or [*BIG*^+^] colonies arising at each timepoint.

### Microscopy and cell size measurements

Most microscopy was performed using a Leica inverted epifluorescence microscope (DMI6000B) with a Hammamatsu Orca 4.0 camera. Cells were imaged after 3 days of growth in 5 mL YPD at 30°C. Saturated cultures were diluted 10-fold with 1X PBS and briefly sonicated to break up cell clumps. Differential interference contrast (DIC) images were taken at 20 millisecond exposure time using a 63x/1.40 oil objective. Cell area was calculated using CellProfiler (3.1.5) image analysis software (www.cellprofiler.org)(Carpenter et al., 2006). Large cell threshold was set at one standard deviation above the cell area mean of the naïve cells.

GFP microscopy data presented in **Figure 4G** was imaged similarly to above, except that cells were grown in YPD for 24 hours.

Cell size experiments using protein synthesis inhibitors (**Fig. 6A–D**): Conditions were the same as above, with the following differences. Single colonies were inoculated into YPD, YPD+cycloheximide (0.05µg/mL), or YPD+rapamycin (10µM) and grown for 4 days before imaging. (Very similar results were observed after 3 days of growth.) The following day, cultures were diluted into liquid YPD and grown for 3 days, after which they were restreaked once onto YPD agar. Single colonies were then used to inoculate liquid YPD cultures, grown for 3 days before imaging to test for the reappearance of the large cell size phenotype.

Data presented in **Figure 8C** was imaged similarly, except that cells were grown in SC-Ura media for retention of the GFP-Pus4 expression plasmid, and imaged using a GE DeltaVision Ultra microscope (Boston, MA).

### Microscopy Image processing

ImageJ version 2.0.0-rc-69/1.52p, Build 269a0ad53f. For GFP-Pus4 in laboratory *S. cerevisiae* (**Fig. 4G**): 1) using full-size images, rolling background subtraction, radius of 50 pixels, 2) enhance contrast to allow 0.1% of pixels to be saturated, 3) check that brightness and contrast are adjusted equivalently in each image.

For GFP-Pus4 in wild *S. cerevisiae* isolate BC187 (**Fig. 8C**): 1) using full-size images, adjust brightness and contrast equivalently in both images, 2) set 50 x 50 pixel square in area between cells and measure average intensity, 3) subtract this intensity from the total image using the “Math” function.

### Luciferase assays

Strains were transformed with PDJ512 and PDJ513. To maintain the plasmids, we grew these cells in a synthetic complete medium containing nutrient levels between those in SD and YPD formulations. Four independent transformants for each sample were grown for 1 day in 150 µL SC-Ura (Sunrise Scientific) per well in 96 well plates at 30°C. Saturated cultures were then diluted 15X into fresh media in a new 96 well plate and grown until cultures reach OD 0.6, as determined by a Biotek Eon plate reader. 20uL of each culture was added using multichannel pipette into a white flatbottom 96-well microplates (E&K Scientific, Santa Clara, CA) already containing 20uL of room temperature 1X Passive Lysis Buffer from the Dual Luciferase Reporter Assay System (Promega, Madison, WI). Cultures were then lysed by shaking at 300 rpm for 25 minutes at room temperature. *Renilla* and firefly luciferase activity was measured using and 75 µL injection volumes and otherwise default settings on a Veritas luminometer (Turner Biosystems). pTH726-CEN-RLuc/minCFLuc (PDJ512) and pTH727-CEN-RLuc/staCFLuc (PDJ513) were gifts from Tobias von der Haar (University of Kent)(Addgene plasmids # 38210 and # 38211) (Chu et al., 2014).

Final luciferase values were normalized to OD measurements of cultures to account for cell density. We note, however, that [*BIG*^+^] cells did not have a general growth advantage over naïve cells in SC-Ura, and when comparing optical density measurements to those counting cells using a hemacytometer, we observed no perturbation in the relationship between cell number and optical density for [*BIG*^+^] cells.

For wild strains, we considered the possibility that curing could reverse multiple epigenetic elements affecting plasmid copy number, transcription, or other elements of gene expression apart from protein synthesis. Indeed, after normalizing *Renilla* or firefly luciferase values to cell density, some strains have several fold-differences after curing, although they were closely correlated irrespective of which firefly codon variant was compared. Therefore, as for data presented in **Figure 6E,** for **Figure 8B** we also normalized firefly luciferase values to *Renilla* luciferase, which is expressed from the same plasmid. This normalization procedure thus tests for differences in protein synthesis that are codon-frequency dependent, i.e. a measure of translational efficiency.

### Polysome profiling

Single colonies from two biological replicates per sample were used to inoculate 5mL YPD cultures that were grown on a roller drum wheel at 30°C overnight. Saturated cultures were added to 95mL of YPD in 500 mL flasks and shaken at 225 rpm at 30°C until cultures reached OD 1.0. Five minutes prior to harvesting cells, we added cycloheximide (Sigma) to final concentration of 100 µg/mL to arrest translation, by adding 1 mL of a 10 mg/mL stock solution (in ethanol) per culture, then immediately swirling flask and putting back on shaker for 5 minutes 225 rpm 30°C to permit the chemical to enter cells and arrest protein synthesis. Cultures were pelleted in 50 mL conical tubes for 3 minutes at 5000 rpm. After decanting supernatant, pellets were quickly resuspend in ice-cold Polysome Lysis Buffer (Jan et al., 2014) (PLB)(20 mM Tris pH 8.0, 140 mM KCl, 1.5 mM MgCl2, 100 µg/mL cycloheximide, 1% Triton X-100, RNase-free reagents), 250 µL total PLB per sample. Resuspended pellets were then flash frozen in liquid nitrogen. Pellets were weighed to ensure their weights were near equal, and then thawed on ice. 250µL of additional ice cold PLB was added per sample, making slightly over 0.5 mL per sample. Samples were then flash frozen in tiny pellets (“yeast dippin’ dots”) by pipetting directly into a small dewar filled with liquid nitrogen and a wire mesh basket nested inside. Tiny pellets were then stored at −80°C until lysis. Samples were lysed using a Retsch Cryomill (Haan, Germany) with 25 mL canisters and the following program: pre-cool, then 12 cycles of 15 Hz x 3 minutes. Smears of lysate were stained with Trypan blue and imaged under a microscope to verify efficient lysis. (We suspect with larger sample volume:canister volume ratios, fewer cycles would be necessary.)

Lysates were loaded onto 10–50% sucrose gradients pre-poured on a BioComp Gradient Master 108 (Fredericton, ND, Canada). Lysates generally contained RNA concentrations around 12–18 µg/µL. 30 µL of lysate was carefully pipetted onto the top of the sucrose gradient, and samples were spun in a Beckman SW41 Ti Rotor for 2.5 hours at 4°C at 40,000 rpm. Gradients were analyzed on Brandel fractionator (Gaithersburg, MD). Technical replicates (same lysate independently loaded onto separate gradients) showed a very high degree of similarity, as did biological replicates.

### GFP-fusion measurements

Naïve or [*BIG*^+^] cells were mated to the SWAT seamless-GFP library (Weill et al., 2018; Yofe et al., 2016) on solid YPD agar plates in 384-spot format for 24 hours at room temperature. Diploids were selected on media lacking both lysine and methionine (SD-Lys-Met) and propagated for 48 hour at room temperature. Diploids were inncoulated into 60 µL of liquid media lacking both lysine and methionine (SD-Lys-Met) in 384-well plates. All library manipulations were carried out using a Singer ROTOR HDA robotic pinning instrument. Cells were propagated in liquid medium for 24 hours at 30°C (OD_600_ ∼ 1), at which time OD_600_ and green fluorescence were measured using a BioTek Synergy H1 plate reader. OD_600_ was adjusted based on known blank wells, and the GFP/OD600 measurements were normalized by Z-score ([*x*_i_ - µ]/σ) within the naïve and [*BIG*^+^] populations independently.

### Western blots

For SDS-PAGE, immunoblots and protein yield measurements, cells were lysed using a Retsch Cryomill using the following program: six 3-minute cycles at 15Hz with 2-minute cooling cycles in between. Cell lysates were loaded onto GenScript ExpressPlus SDS-PAGE 4–20% gels (Piscataway, NJ) and stained using coomassie blue. For Western Blots, anti-FLAG M2 monoclonal antibody (Sigma) was used to detect 3XFLAG-Pus4, and anti-PGK1 monoclonal antibody (Invitrogen) was used to detect the loading control.

### RNA-sequencing

Five independent colonies of naïve and [*BIG*^+^] cells were grown in 5 mL YPD cultures on a culture roller drum wheel overnight at 30°C. Saturated cultures were added to 700 mL YPD cultures in 2 L narrow mouthed baffled flasks, pre-incubated to 30°C. These were grown until a cuvette reading of OD 1.0. (700 mL cultures were used to provide enough material for other sample-intensive experiments done in parallel). Cultures were pelleted in large table-top centrifuge with swing-bucket rotor at 4300 x g at 4°C for 20 minutes, washing once with ice-cold 1X TBS. RNA was harvested as described in (Carlile et al., 2015). Total RNA was treated with RiboZero (Illumina, San Diego, CA) to remove most rRNAs but retain mRNAs and other ncRNAs. RNA was fragmented and submitted for Illumina sequencing at the Beijing Genomics Institute. Analysis was performed using Bowtie2 (Langmead and Salzberg, 2012), HT-Seq (Anders et al., 2015), and DE-Seq2 (Love et al., 2014). Raw data and other experimental information are available on the Gene Expression Omnibus (Barrett et al., 2013), accession: https://www.ncbi.nlm.nih.gov/geo/query/acc.cgi?acc=GSE153930

### tRNA/rRNA quantification

Strains were grown up in 5mL of YPD overnight at 30°C. Saturated cultures were then diluted 15X into fresh media and grown up to an OD 0.8. Cells were then pelleted, and total RNA was extracted by phenol-chloroform extraction. RNA concentration for each sample was then analyzed on Nanodrop, and diluted to equal concentrations in RNase-free water. The extracted RNA was then run on nucleic acid fragment analyzer, and total tRNA abundance was quantified relative to abundance of 5.8S rRNA.

### Pseudouridine measurements

Protocol adapted from reference (Lei and Yi, 2017). For each sample, 40 µg of total RNA was fragmented in RNA fragmentation buffer (New England Bio Labs, Ipswich, MA) at 94°C for 3 min. Following ethanol precipitation, fragmented RNA was resuspended in 80 µL of 5 mM EDTA, denatured at 80°C for 5 min, and then immediately chilled on ice. Each sample was then split into two 40 µL samples for a CMC-labeled and non-labeled control. The 40 µL RNA sample destined for CMC-labeling was added to 400 µL BEU+CMC buffer (50 mM Bicine, pH 8.5; 4 mM EDTA; 7 M urea; 200 mM CMC (Sigma)). The non-labeled 40 µL RNA sample was added to 400µL BEu buffer (50 mM Bicine, pH 8.5; 4 mM EDTA; 7 M urea). Both samples were incubated at 37°C for 20 min to carry out the CMC-Ψ reaction, followed by an ethanol precipitation. Each sample was then resuspended in 200 µL Na_2_CO_3_ buffer (50 mM Na_2_CO_3_, pH 10.4; 2 mM EDTA) and incubated at 37°C for 6 h. Following incubation, RNA was ethanol precipitated and resuspended in 40 µL H2O. RNA was annealed to primers by the addition of 4 µL 100 µM Random Hexamer Primers (TaKaRa, Mountain View, CA) and incubation at 65°C for 5 min. Samples were chilled on ice afterwards. To perform the reverse transcription, 32 µL RT Buffer (125 mM Tris, pH 8.0; 15 mM MnCl_2_; 187.5 mM KCl; 1.25 mM dNTPs; 25mM DTT) was added to each sample. Samples were then incubated at 25°C for 2 min. After, 0.5 µL SuperScript II reverse transcriptase (SSII, Invitrogen, Waltham, MA) was added to each sample followed by incubation at 25°C for 10 min, 42°C for 3 h, and 70°C for 15 min. To perform qPCR analysis, 2 µL of sample was mixed with 10 µL 2X SYBR Mix (Kapa Biosystems, Wilmington, MA), 0.4 µL of each 10 µM primer (IDT, Coralville, IA), and 7.2 µL H2O for a total of 20 µL for each reaction. qPCR was performed in a Bio-Rad CFX Connect Real-Time System (Bio-Rad, Hercules, CA) using the following protocol: initial incubation at 95°C for 5 min, followed by 45 cycles of 95°C for 0.5 min and 60°C for 1 min. Following amplification, reaction was brought down to 54°C and held for 5s, increasing in temperature by 0.1°C increments until 95°C is reached to obtain melt curve data.

### Data display

Plots/graphs were made using PRISM 7/8 software (GraphPad, San Diego, CA).

